# Trait-Relevant Tasks Improve Personality Prediction from Structural-Functional Brain Network Coupling

**DOI:** 10.1101/2025.09.17.676801

**Authors:** Johanna L. Popp, Jonas A. Thiele, Joshua Faskowitz, Caio Seguin, Olaf Sporns, Kirsten Hilger

## Abstract

Personality traits capture stable patterns of behavior and thought, and neurobiological correlates were identified in structural and functional brain networks. Here, we investigate whether the coupling between structural and functional brain networks (SC-FC coupling), during resting state and seven tasks of varying trait-relevance, is associated with individual differences in the Big Five personality traits. We used diffusion-weighted and functional magnetic resonance imaging from 764 participants of the Human Connectome Project and modelled individual differences in SC-FC coupling with similarity and communication measures. These measures approximate functional interactions unfolding on top of the structural connectome and were set in relation to individual variations in personality traits. Significant associations were only observed during trait-relevant tasks: for agreeableness during social cognition, and conscientiousness could be predicted from task-general coupling patterns. We conclude that optimizing trait-relevance of tasks during neuroscientific measurements presents a promising means to increase effect sizes in studies on brain-behavior associations.

## 1. Introduction

Individuals differ in personality, defined as a unique and consistent pattern of thought, emotion, and behavior. Personality traits represent core dimensions of this pattern (DeYoung, 2010, 2015) and individual differences in traits have been associated with vulnerability for psychopathology (Kotov et al., 2010) and critical life outcomes such as academic and occupational success (Poropat, 2009; Rothmann & Coetzer, 2003). Understanding their neurobiological basis therefore presents a highly relevant research topic.

### 1.1. The Big Five personality traits

The five-factor model of personality (Costa & McCrae, 1992b; Goldberg, 1990) is the most widely used taxonomy of personality traits. Although it was initially developed through the analysis of trait-descriptive terms in English dictionaries (Allport & Odbert, 1936; Cattell, 1945), the Big Five personality traits – agreeableness (A), openness to experience (O), conscientiousness (C), neuroticism (N), and extraversion (E) – have been shown to emerge across diverse cultures, suggesting their universality. The manifestation of individual differences in personality traits in overt behavior was assumed to depend on situational context, particularly on situational trait-relevance (e.g., individual extraversion levels are better observable during parties than during lectures; DeYoung, 2015; Hilger & Markett, 2021; Tett & Burnett, 2003; Tett & Guterman, 2000). However, whether the trait-relevance of tasks enhances the detectability of neural correlates of personality traits remains an open question.

### 1.2. Neural correlates

Genetic influences on variations in the Big Five (40-80% heritability; DeYoung, 2010) suggest their biological foundation. In line with this, between-person differences have been associated with region-specific characteristics of brain structure and function (Canli, 2004; DeYoung, 2010; DeYoung et al., 2010, 2021). However, such findings were inconsistent and sometimes contradictory (Y.-W. Chen & Canli, 2022; Toschi et al., 2018). Network neuroscience, conceptualizing the human brain as a complex network, emerged as promising framework to resolve this heterogeneity (Bassett & Sporns, 2017; Bullmore & Sporns, 2009). Variations in complex human traits, like the Big Five, were proposed to arise from brain-wide interactions rather than from attributes of individual regions (Hilger & Markett, 2021; Liu et al., 2019). Accordingly, characteristics of structural connectivity (SC; tractography performed on diffusion-weighted imaging [DWI]) and functional connectivity (FC; statistical dependencies of blood-oxygen-level-dependent [BOLD] signal time courses) have been related to individual differences in the Big Five (e.g., Dubois et al., 2018; Nostro et al., 2018; Ueda et al., 2018). However, whether the interplay of both modalities, the structural-functional brain network coupling (SC-FC coupling), relates to differences in personality traits remains unclear.

### 1.3. SC-FC coupling

Different methods have been developed to capture this interplay between structure and function. Complex biophysical models, incorporating plausible biological mechanisms to link SC and FC, are often impractical given their high computational costs (Murray et al., 2018; Suárez et al., 2020). In contrast, statistical approaches, e.g., correlational analyses, lack explanatory power regarding underlying biological mechanisms. As a compromise, the operationalization of SC-FC coupling with similarity and communication measures offers an efficient yet informative approach with proven potential for individual differences research (Popp et al., 2024, 2025; Seguin et al., 2020; Zamani Esfahlani et al., 2022). While similarity measures assess the similarity of regional structural connectivity profiles and thus serve as baseline for SC-FC coupling when compared to FC, communication measures approximate FC by quantifying the ease of communication between two brain regions based on SC. Communication processes are estimated according to theoretical network modeling, encompassing regimes from routing to diffusion (Avena-Koenigsberger et al., 2018; Goñi et al., 2014; Seguin, Sporns, et al., 2023; Suárez et al., 2020). The extent to which these communication measures align with the observed FC could be informative of underlying neural signaling processes (Goñi et al., 2014; Seguin, Jedynak, et al., 2023) and thus offers insights into biological mechanisms underlying behavioral variability. However, this approach has not yet been applied in the investigation of personality trait differences.

### 1.4. The current study

This preregistered study investigates the relationship between individual differences in the Big Five personality traits and variations in SC-FC coupling. To this aim, we used neuroimaging data from 764 participants of the Human Connectome Project (HCP; Van Essen et al., 2013) and operationalized SC-FC coupling with one similarity measure and three communication measures. Neuroimaging data was acquired during resting state and seven tasks differing in their trait-relevance. We examined correlative associations at the brain-average level, explored potential influences of tasks’ trait-relevance, and developed a predictive modelling framework to ultimately predict individual personality trait scores. All analyses were validated in a held-out lockbox sample.

## 2. Methods

### 2.1. Transparency and Openness

We report how we determined our sample size, all data exclusions, all manipulations, and all measures in the study. The code for all analyses, including preprocessing, is available on GitHub: DWI preprocessing: https://github.com/civier/HCP-dMRI-connectome; Functional magnetic resonance imaging (fMRI) preprocessing: https://github.com/faskowit/app-fmri-2-mat; Computation of latent *g*-factor: https://github.com/jonasAthiele/BrainReconfiguration_Intelligence; Operationalization of SC-FC coupling: https://github.com/brain-networks/local_scfc; Main analysis and validation analysis: https://github.com/johannaleapopp/SC_FC_Coupling_Task_Personality. Data from the Human Connectome Project can be downloaded online under https://www.humanconnectome.org/study/hcp-young-adult (Human Connectome Project Consortium, 2017). MATLAB code (version 2021a) for all here reported analyses has been deposited on Zenodo (https://doi.org/10.5281/zenodo.17131036; Popp, 2025).

### 2.2. Preregistration

This study is part of a larger research project focusing on the relationships between individual differences in complex traits and SC-FC coupling. The complete project has been preregistered in the Open Science Framework (https://osf.io/ctvf9), encompassing plans for the current study and additional studies focused on intelligence (Popp et al., 2025).

### 2.3. Participants

All analyses were conducted on the HCP Young Adult Sample S1200 (Van Essen et al., 2013), comprising 1,200 participants (656 female; 1089 right-handed; mean age = 28.8 years; age range = 22-37 years).

We excluded participants with a) unavailable DWI, resting-state or task fMRI, b) missing personality or cognitive scores required to compute a latent intelligence factor (control analyses), and c) Mini-Mental State Examination scores < 27, indicating cognitive impairment. Further, participants who met at least one criterion for excessive head motion (Parkes et al., 2018) during any fMRI condition, as measured by framewise displacement (FD; Jenkinson et al., 2002), were excluded. These criteria were: mean FD > 0.20 mm; proportion of spikes (FD > 0.25 mm) > 20 percent; any motion spikes > 5.00 mm.

The final sample comprised 764 participants (402 female; 697 right-handed; mean age = 28.6 years; age range = 22-36 years) and was split 70/30 before analysis into a main sample (HCP532; *N* = 532; 271 female; 485 right-handed; mean age = 28.6 years; age range 22-36 years) and a lockbox sample (HCP232; *N* = 232; 131 female; 212 right-handed; mean age = 28.5 years; age range = 22-36 years) retained for validation.

### 2.4. Personality traits

The Big Five personality traits (Table 1) were assessed with the NEO-Five-Factor Inventory (NEO-FFI), a 60-item version of the Costa and McCrae Five-Factor Inventory (Costa & McCrae, 1992a; McCrae & Costa Jr., 2004). Participants rated their agreement with 12 trait-specific statements reflecting their self-perception (e.g., I am not easily disturbed) on a five-point Likert scale (strongly disagree = 0; strongly agree = 4).

**Table 1.** The Big Five personality traits.

| Personality trait | Description |
| --- | --- |
| Agreeableness (A) | Interpersonal behavior; degree of avoiding interpersonal conflicts; tendency towards altruism as opposed to exploitation of others. |
| Openness to experience (O) | Tendency to engage with aesthetic, intellectual and sensory information; degree of appreciation/toleration of the unfamiliar. |
| Conscientiousness (C) | Degree of self-discipline, organization and aim for achievement; ability to control behavior and impulses in order to follow rules and to pursue non-immediate goals. |
| Neuroticism (N) | Sensitivity to punishment and negative emotions; tendency to experience psychological distress. |
| Extraversion (E) | Sensitivity to reward and positive affect; approach behavior; tendency to experience positive emotions such as joy and pleasure. |

*Note:* Descriptions were adapted from DeYoung (2010) and Nostro et al. (2018).

### 2.5. Neuroimaging data acquisition and preprocessing

MRI scans were acquired using a Siemens Skyra 3T scanner with a 32-channel head coil. Details are described below and in Van Essen et al. (2013).

#### 2.5.1. Diffusion-weighted imaging

SC was estimated based on the HCP’s minimally preprocessed DWI data (time to repetition (TR) = 5520 milliseconds (ms); time to echo (TE) = 89.5 ms; 1.25 mm isotropic voxel resolution; multiband acceleration factor = 3; b = 1000, 2000, 3000 s/mm^2^; 90 directions/shell; Glasser et al., 2013). Following the MRTrix pipeline (Civier et al., 2019; Tournier et al., 2019), bias correction, white matter fiber modeling via constrained spherical deconvolution (Tournier et al., 2007), and tissue normalization (Dhollander et al., 2021) were implemented. Finally, probabilistic streamline tractography was applied (R. E. Smith et al., 2012), retaining only streamlines aligning with the estimated white matter orientations from the diffusion image (Glasser et al., 2013; R. E. Smith et al., 2013; Tournier et al., 2012).

#### 2.5.2. Functional magnetic resonance imaging

FMRI scans were acquired in two sessions using a gradient-echo EPI sequence with multi-slice acceleration (TR = 720 ms; TE = 33.1 ms; 2-mm isotropic voxel resolution; flip angle = 52°; multiband acceleration factor = 8). Each session included two 15-minute resting-state scans in opposing phase encoding directions (L/R and R/L) followed by approximately 30 minutes of task fMRI, consisting of seven tasks that were divided across the two sessions (eight total conditions, Table 2; each L/R and R/L). After downloading the minimally preprocessed fMRI data, 24 motion parameters, eight mean signals from white matter and cerebrospinal fluid, and four global signals were regressed out (Parkes et al., 2018), while task-evoked activation was removed with basis-set task regressors (Cole et al., 2019).

**Table 2.** Trait-relevance of fMRI conditions.

|  | Condition | Description | Frames per run | Run duration (min) | Trait-relevance |
| --- | --- | --- | --- | --- | --- |
| 1 | Resting state (RES) | Participants keep eyes open and focus on a fixation cross. | 4*1200 | 14:33 | - |
| 2 | Emotion processing (EMO) | Matching of angry and fearful faces or shapes. | 2*176 | 2:16 | A |
| 3 | Gambling (GAM) | Card guessing game with possibility of winning or losing money. | 2*253 | 3:12 | C, N, E |
| 4 | Language (LAN) | Auditory presentation of a brief story, followed by a question related to the story's topic. | 2*316 | 3:57 | O |
| 5 | Motor (MOT) | Visual cues instructing participants to tap their fingers, squeeze their toes or move their tongue. | 2*284 | 3:34 | - |
| 6 | Relational processing (REL) | Presentation of two pairs of objects, followed by a decision whether the objects have similar shapes or textures. | 2*232 | 2:56 | O |
| 7 | Social cognition (SOC) | Presentation of short videos featuring interacting versus randomly moving objects (squares, circles, triangles), followed by a decision on whether a social interaction had occurred. | 2*274 | 3:27 | A, E |
| 8 | Working memory (WM) | 2-back and 0-back working memory task involving faces, places, tools, and body parts. | 2*405 | 5:01 | O |

*Note:* Resting-state and task fMRI data were acquired in two sessions. Scan length differed between fMRI conditions, but image acquisition parameters did not (Barch et al., 2013). Trait-relevance of fMRI tasks was determined based on existing literature and preregistered. A = Agreeableness; O = Openness to experience; C = Conscientiousness; N = Neuroticism; E = Extraversion. Table adapted from Popp et al. (2025).

### 2.6. Trait-relevance of tasks

Trait-relevance of fMRI tasks was determined based on existing literature and preregistered (Table 2, details in Supplemental Material p. 41).

### 2.7. Structural and functional brain network connectivity

To construct individual structural and functional brain networks, Glasser et al.’s multimodal parcellation scheme (2016), dividing the cortex into 360 brain regions, was used. As left and right hippocampi were classified as subcortical during preprocessing, 358 regions were included. Individual SC matrices entail SIFT2 streamline density weights between pairs of brain regions, while individual FC matrices for each fMRI condition comprise Fisher-*z* transformed Pearson correlations of regional BOLD signal time courses.

SC matrices were modelled as symmetric and condition-specific FC matrices were computed separately for each phase encoding direction and averaged afterwards (Cole et al., 2014; Popp et al., 2024; S. M. Smith et al., 2013).

### 2.8. Structural-functional brain network coupling

Building on each individuals’ SC, we computed one similarity measure (cosine similarity, CoS) and three communication measures (path length, PL; communicability, G; search information, SI), each retaining the same matrix dimensions (358 x 358; Figure 1). These communication measures were selected to capture the entire spectrum of communication models from routing to diffusion (Seguin, Jedynak, et al., 2023; Seguin, Sporns, et al., 2023; for details: Table 3; Supplemental Material pp. 42-44). To compute brain region-specific SC-FC coupling values, regional connectivity profiles (matrix columns containing information about the connections one specific brain region has to all other regions) of each similarity or communication matrix were correlated with the corresponding regional connectivity profile of a condition-specific FC matrix (4 similarity/communication measures * 8 fMRI conditions = 32 comparisons).

**Figure 1.**
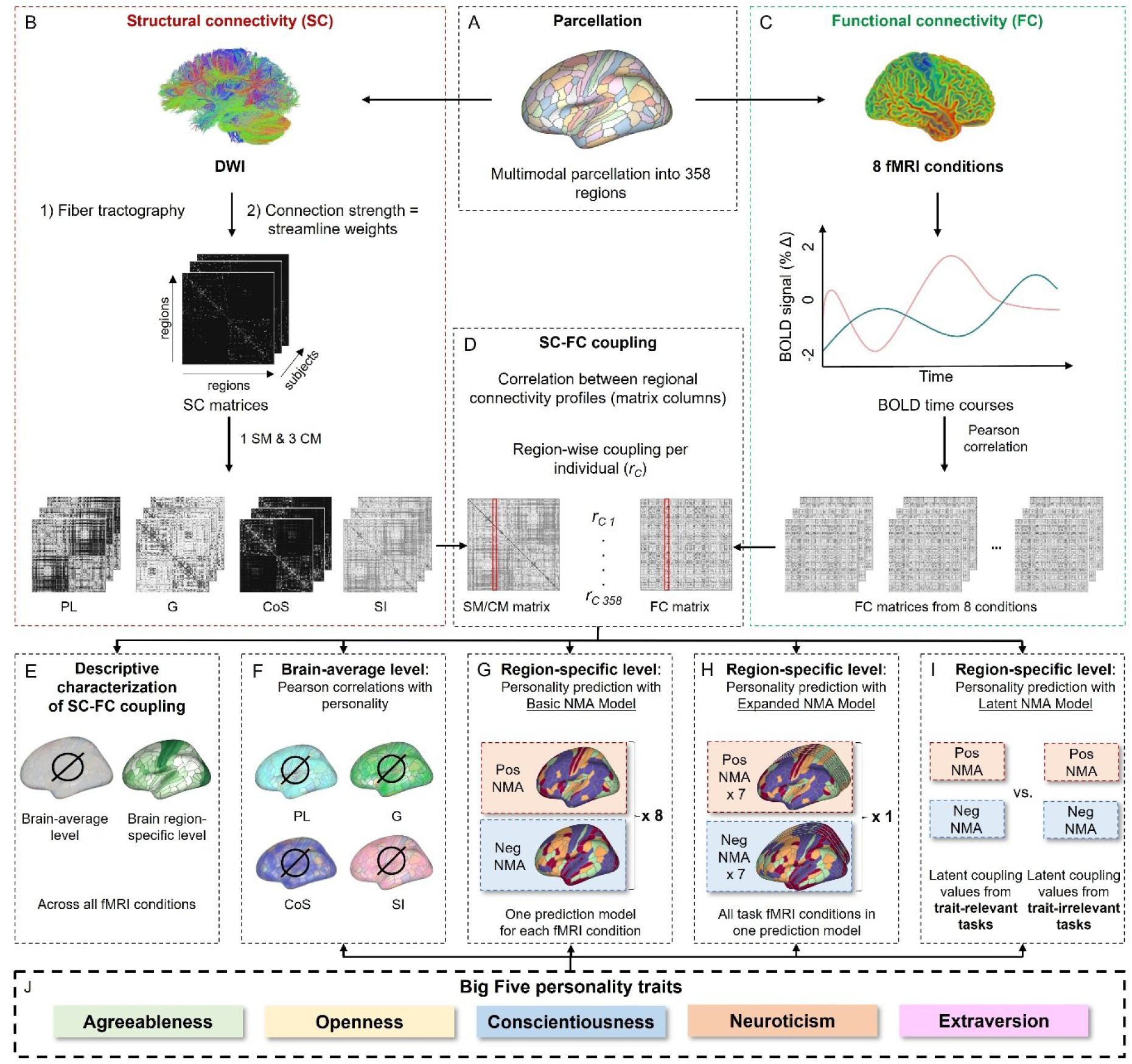
Schematic overview of study procedure. (A) DWI and fMRI data were parcellated into 358 brain regions based on a multimodal parcellation scheme by Glasser et al. (2016). (B) Individual structural connectivity matrices were transformed into one similarity matrix, capturing similarity in regional SC, and three communication matrices, approximating functional interactions on top of the underlying structural connectome under an assumed model of network communication (Table 3). (C) For each of the eight fMRI conditions (Table 2), one individual FC matrix was constructed by performing Pearson correlations between all possible pairs of regional BOLD signal time courses. (D) To compute region-specific SC-FC coupling values, each individual’s similarity matrix and three communication matrices were compared to the condition-specific FC matrices (4*8 comparisons). Specifically, regional connectivity profiles (matrix columns) of the respective similarity or communication matrix were correlated with the corresponding regional connectivity profile of the condition-specific FC matrix. For each fMRI condition, four comparisons were made. Each yielded 358 individual SC-FC coupling values (*r_c_*, one per brain region), which constitute the four condition-specific coupling measures. (E) Investigation of group-general differences in SC-FC coupling (see Popp et al., 2025). (F) Relationship between personality traits and brain-average SC-FC coupling. (G) Prediction of individual personality trait scores from region-specific SC-FC coupling with the B-NMA using data from one condition at a time. A more detailed illustration and description of this prediction model can be found in Figure S1. (H) Prediction of personality traits from region-specific SC-FC coupling with the E-NMA integrating data from all seven tasks at a time. A more detailed illustration and description of this prediction model can be found in Figure S2. (I) Prediction of personality traits from region-specific SC-FC coupling, contrasting trait-relevant versus trait-irrelevant latent coupling factors with the L-NMA. A more detailed illustration and description of this prediction model can be found in Figure S3. (J) Big Five personality traits were measured with the NEO-FFI (Costa & McCrae, 1992a; McCrae & Costa Jr., 2004). SC = Structural brain network connectivity; FC = Functional brain network connectivity; BOLD = Blood oxygen level dependent; SM = Similarity measure; CM = Communication measure; PL = Path length; G = Communicability; CoS = Cosine similarity; SI = Search information; NMA = Node-measure assignment. Figure adapted from Popp et al. (2025).

**Table 3.** Measures of SC-FC coupling.

| Similarity measure | Description | Reference |
| --- | --- | --- |
| Cosine similarity (CoS) | Similarity of connectivity profiles based on vector orientation. | (Han et al., 2012) |
| **Network communication measure** |  |  |
| Path length (PL) * | Length of the shortest possible path between brain regions. | (Avena-Koenigsberger et al., 2018) |
| Communicability (G) | Communication via diffusive broadcasting process. In a network, all walks (sequence of traversed edges) of all lengths are considered and the contribution of each walk is inversely proportional to its length. | (Estrada & Hatano, 2008) |
| Search information (SI) ** | Probability that a random walker will travel between two nodes via their shortest path. | (Rosvall et al., 2005) |

*Note:* Measures whose matrices were adjusted after computation are marked with an asterisk (\* = matrix was inverted; \*\* = matrix was symmetrized and inverted, Supplemental Material pp. 42-44; Table adapted from Popp et al. (2025).

### 2.9. Descriptives of structural-functional brain network coupling

#### 2.9.1. Brain-average level

Per participant, brain-average SC-FC coupling values were computed for each combination of fMRI condition (Table 2) and coupling measure (CoS, PL, G, SI) by averaging the corresponding 358 region-specific SC-FC coupling values (*r*_c_).

#### 2.9.2. Brain region-specific level

For visualizing differences in regional SC-FC coupling patterns between conditions, we first determined, per condition and brain region separately, which similarity or communication measure explained the highest variance in FC across participants. This condition-specific group-general mapping between brain regions and coupling measures was then used to extract individual region-specific SC-FC coupling values. These were then averaged across all participants to create a vector with 358 elements, defining the group-general map of condition-specific SC-FC coupling.

### 2.10. Structural-functional brain network coupling and personality traits

#### 2.10.1. Associations with personality traits: Brain-average level

Partial Pearson correlations between individual personality trait scores and individual brain-average SC-FC coupling values were computed (separately for each combination of fMRI condition and coupling measure), while controlling for age, gender, handedness, and in-scanner motion (condition-specific mean FD). Significance was determined using a Bonferroni-corrected threshold for four comparisons: *p* < .0125.

#### 2.10.2. Prediction of personality: Brain region-specific level

##### 2.10.2.1. Basic node-measure assignment model (B-NMA): Personality trait prediction from SC-FC coupling during single fMRI conditions

The B-NMA, a current regression-based prediction framework (Popp et al., 2024, 2025), was applied to test whether individual personality traits can be predicted from brain region-specific SC-FC coupling. Predictions were performed separately for each personality trait, while using input from one single fMRI condition and applying 5-fold cross-validation. Concisely, per condition, scores for the trait of interest from participants in the training sample were correlated with region-specific SC-FC coupling values (*r_C_*; partial correlations controlling for effects of age, gender, handedness, and motion). This was done for each coupling measure separately, resulting in four correlation coefficients (*r_P_*) per brain region. Next, two training group-general node measure assignment (NMA) masks were generated by assigning the coupling measure with the largest positive magnitude association (positive NMA) and the measure with the largest negative magnitude association (negative NMA) between personality trait score and condition-specific SC-FC coupling values (*r_C_*) to a given brain region. These two masks were used to extract individual region-specific SC-FC coupling values (*r_C_*) and two model input features were created: one by averaging across all individual condition-specific SC-FC coupling values extracted with the positive NMA mask, and the other by averaging across all values extracted with the negative NMA mask. Importantly, NMAs from the training samples were always used to extract region-specific SC-FC coupling values in the test samples. For complementary illustrations and detailed descriptions see Figure S1 and Popp et al. (2024).

##### 2.10.2.2. Expanded node-measure assignment model (E-NMA): Personality trait prediction from SC-FC coupling combined across all tasks

The E-NMA was developed to assess whether the integration of SC-FC coupling information across tasks improves personality prediction. Also here, two model input features were generated for each fMRI condition (excluding resting state), but the E-NMA simultaneously incorporates all 14 condition-specific features into the regression model (Figure S2).

##### 2.10.2.3. Latent node-measure assignment model (L-NMA): Personality trait prediction from SC-FC coupling during trait-relevant versus trait-irrelevant tasks

The L-NMA contrasts prediction of personality traits from region-specific SC-FC coupling during trait-relevant versus trait-irrelevant tasks. Here, two separate prediction models were created for each personality trait by combining region-specific SC-FC coupling information from a) all trait-relevant tasks and b) all trait-irrelevant tasks into latent SC-FC coupling values, before creating the two model input features with the same strategy as in the B-NMA. Crucially, these latent SC-FC coupling values were either computed as a latent factor (factor 1; if more than two tasks were trait-relevant/trait-irrelevant) or as the first component of a principal component analysis (PCA; if two tasks were trait-relevant/trait-irrelevant; Figure S3). In case only one trait-relevant/trait-irrelevant task existed, the B-NMA was used.

##### 2.10.2.4. Prediction performance

Prediction performance was evaluated via Pearson correlation between predicted and observed personality trait scores. For robustness, 100 repetitions with varying training-test splits were performed and prediction accuracy was averaged. Statistical significance was assessed with non-parametric permutation tests (1000 iterations). Significant associations uncorrected for multiple comparisons (*p* < .05) are presented in the Supplemental Material, while those meeting the Bonferroni-corrected threshold (eleven comparisons, *p* < .005) are reported in the main text.

### 2.11. Lockbox validation

To evaluate the robustness and generalizability of our findings, all analyses were repeated in the lockbox sample. If possible, additional cross-sample model generalization tests were performed: The main sample data were split into five folds and five prediction models were trained, each on 80% of the data (4/5 folds). These models were then used to predict individual personality trait scores in the lockbox sample. This procedure was conducted 100 times with different data splits. The performance of each test was evaluated by averaging across all prediction performances, measured as Pearson correlations between predicted and observed scores. Model significance was determined by comparing prediction performances when the model was trained on observed versus permuted personality trait scores (1000 iterations).

## 3. Results

### 3.1. Personality traits

Individual agreeableness scores ranged from 13 – 46 (*M* = 33.8; *SD* = 5.7), openness to experience from 13 – 47 (*M* = 28.7; *SD* = 6.2), conscientiousness from 11 – 48 (*M* = 34.3; *SD* = 5.7), neuroticism from 1 – 41 (*M* = 16.5; *SD* = 7.47) and extraversion from 10 – 47 (*M* = 30.2; *SD* = 6.3; frequency distributions in Figure S4; inter-correlations in Table S1).

### 3.2. Descriptives of SC-FC coupling

As detailed characterizations of brain-average and brain region-specific SC-FC coupling, including measure- and condition-specific differences, are reported in Popp et al. (2025), only relevant aspects for the subsequent analyses are listed below.

#### 3.2.1. Brain-average level

To visualize condition-specific brain-average SC-FC coupling, all four measure-specific SC-FC coupling values (*r_C_*) were averaged per participant (Figure S5A). Likewise, for measure-specific brain-average SC-FC coupling, we averaged the eight condition-specific SC-FC coupling values (*r_C_*) per participant (Figure S5B). Brain-average SC-FC coupling was highest during resting state and lowest during the emotion processing task, while brain-average SC-FC coupling was highest when operationalized with cosine similarity and lowest when measured with search information.

#### 3.2.2. Brain region-specific level

Across all conditions, the group-average region-specific SC-FC coupling pattern was characterized by higher coupling in unimodal areas and lower coupling in multimodal areas (Figure S5C). SC-FC coupling patterns across different tasks were highly similar, whereas resting-state SC-FC coupling showed greater divergence.

### 3.3. Personality traits and structural-functional brain network coupling

#### 3.3.1. Unconstrained state: No relationship between SC-FC coupling and personality traits during resting state

Personality traits were not significantly associated with measure-specific brain-average SC-FC coupling during resting state (Table S2). Similarly, no personality trait could be significantly predicted from region-specific SC-FC coupling during resting state using the B-NMA (all *r* ≤ 0.07; all *p* > .05; Table S3).

#### 3.3.2. Facing tasks: Specific relationships with personality traits emerge when constraining the situational context

Agreeableness was significantly positively associated with brain-average SC-FC coupling during the social cognition task when operationalized with search information (*r* = .12; *p* = .006; Figure 2A). All other associations did not reach significance (all *r* ≤ .09, all *p* > .0125; Tables S4-S10).

**Figure 2.**
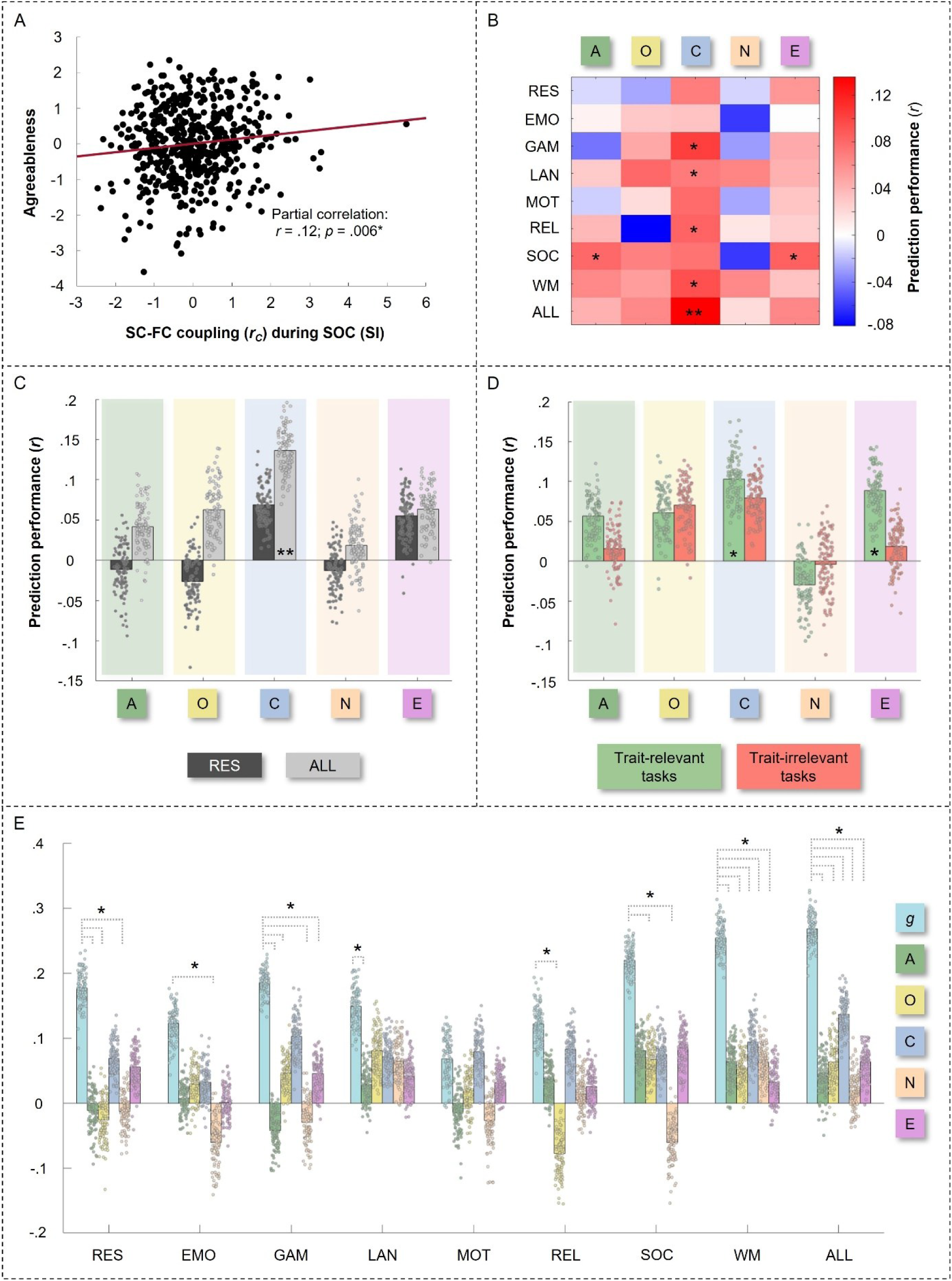
The Big Five personality traits and structural-functional brain network coupling. (A) Positive association between agreeableness and brain-average SC-FC coupling during the social cognition task (Table S9). (B) Performance when predicting personality traits with brain region-specific SC-FC coupling from single tasks using the B-NMA (Table S3). (C) Performance of personality trait prediction from task-combined region-specific SC-FC coupling with the E-NMA versus from resting-state region-specific SC-FC coupling with the B-NMA: Task-combined SC-FC coupling descriptively outperforms resting-state SC-FC coupling for all personality traits while only conscientiousness can be significantly predicted from task-combined SC-FC coupling (Table S11). (D) Performance of personality trait prediction from latent region-specific SC-FC coupling values during trait-relevant versus trait-irrelevant tasks with the L-NMA: No significant influence of trait-relevance (Table S12). (E) Post-hoc analysis: Better prediction of general intelligence than personality traits from region-specific SC-FC coupling with the B-NMA and E-NMA (Table S25-S27). Significant predictions (Figure 2B-D) and model difference tests (Figure 2E) uncorrected for multiple comparisons are marked with one asterisk (* = *p* < .05, reported in the Supplemental Material), while significant predictions passing the Bonferroni-corrected threshold (for eleven comparisons; Figure 2B-D) are marked with two asterisks (** = *p* < .005). RES = Resting state; EMO = Emotion processing task; GAM = Gambling task; LAN = Language task; MOT = Motor task; REL = Relational processing task; SOC = Social cognition task; WM = Working memory task; ALL = All task conditions (i.e. prediction with E-NMA); A = Agreeableness; O = Openness to experience; C = Conscientiousness; N = Neuroticism; E = Extraversion; *g* = General intelligence.

Only conscientiousness could be significantly predicted from task-combined region-specific SC-FC coupling (E-NMA model; *r* = .14; *p* = .001; Figure 3, Table S11). However, descriptively, prediction from task-combined region-specific SC-FC coupling (E-NMA) outperformed prediction from region-specific SC-FC coupling during resting state for all personality traits (B-NMA; Figure 2B-2C; Table S11).

**Figure 3.**
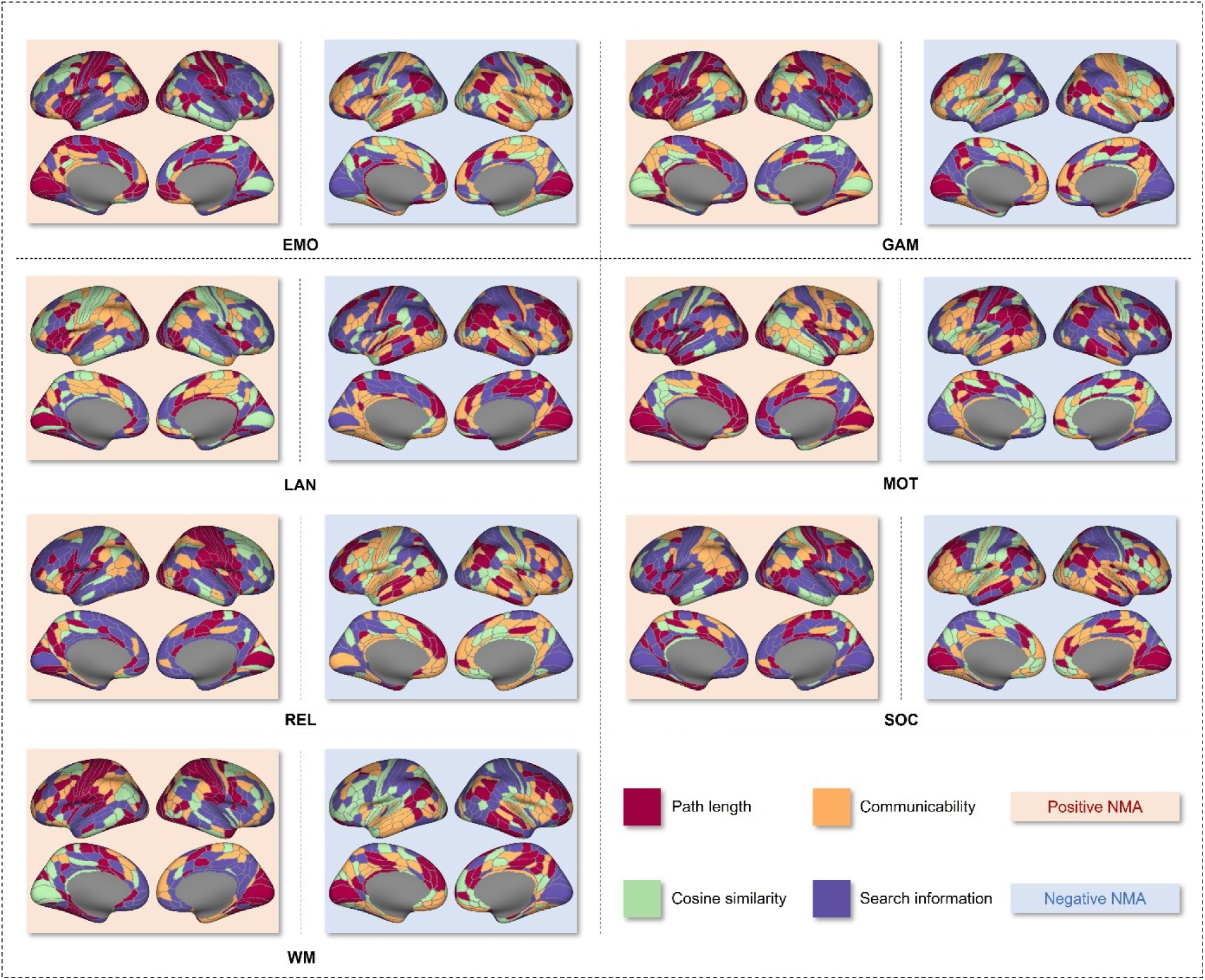
Conscientiousness can be predicted from task-combined brain-region specific structural-functional brain network coupling. Assignments of coupling measures to brain regions (NMAs) were based on the largest positive (red background) and largest negative (blue background) magnitude associations between conscientiousness scores and measure-specific SC-FC coupling values per brain region across all participants. These masks were generated separately for each condition and consequently used to extract individual region-specific SC-FC coupling values, which were then averaged – separately for each of the fourteen cases – to serve as input features for the significant prediction of individual conscientiousness scores (*r* = .14; *p* = .001). Note that the visualized NMAs were created in the complete main sample, while the NMAs actually applied in the prediction models were created only based on data from the training samples, thus avoiding any data leakage. RES = Resting state; EMO = Emotion processing task; GAM = Gambling task; LAN = Language task; MOT = Motor task; REL = Relational processing task; SOC = Social cognition task; WM = Working memory task.

Overall, prediction performance did not differ between trait-relevant and trait-irrelevant tasks when predicting from region-specific SC-FC coupling. This holds true when predicting personality traits from region-specific SC-FC coupling with the B-NMA (Figure 2B; Table S3), as well as when comparing prediction performance between latent brain region-specific SC-FC coupling values from all trait-relevant versus all trait-irrelevant tasks with the L-NMA (Figure S2D; Table S3; Table S12).

### 3.4. Validation

#### 3.4.1. Personality traits

In the lockbox sample, agreeableness scores ranged from 15 – 47 (*M* = 33.9; *SD* = 5.8), openness to experience from 10 – 44 (*M* = 28.8; *SD* = 6.3), conscientiousness from 21 – 48 (*M* = 34.9; *SD* = 5.7), neuroticism from 0 – 39 (*M* = 15.7; *SD* = 6.9) and extraversion from 14 – 45 (*M* = 31.5; *SD* = 5.3; Frequency distributions: Figure S6).

#### 3.4.2. Descriptives of SC-FC coupling

Similar to the main sample, brain-average SC-FC coupling was highest during resting state and lowest during the emotion processing task. Again, highest brain-average SC-FC coupling was observed when operationalized with cosine similarity, while lowest coupling resulted from search information. Also the group-average cortex-wide distribution of SC-FC coupling strength was similar to the main sample: higher coupling in unimodal areas and lower coupling in multimodal areas (Figure S7; Popp et al., 2025).

#### 3.4.3. Personality traits and structural-functional brain network coupling

Contrasting the main analyses, one association between personality traits and resting-state brain-average SC-FC coupling passed the corrected significance threshold (openness to experience & coupling via search information: *r* = .17; *p* = .009; Table S13). However, like in the main sample, no personality trait could be significantly predicted from region-specific SC-FC coupling during resting state, neither by using the 5-fold internally cross-validated B-NMA (all *r* ≤ .08; all *p* > .005; Table S14) nor by using the cross-sample model generalization test applying the B-NMA with parameters derived from the main sample on the lockbox sample (all *r* ≤ .06; all *p* > .005; Table S22).

Regarding task-induced coupling, there was no significant association between any personality trait and brain-average SC-FC coupling (all *p* > .0125; Tables S15-S21). Like in the main sample, personality trait predictions from task-combined SC-FC coupling (E-NMA) descriptively outperformed predictions from resting-state and single task region-specific SC-FC coupling (B-NMA; 5-fold internal cross-validation framework; Table S23). This was not true for the cross-sample model generalization test (Table S24). The prediction of conscientiousness from task-combined SC-FC coupling (E-NMA) was neither significant in the lockbox sample (*r* = .08; *p* = .142; Table S23), nor in the cross-sample model generalization test (*r* = −.01; *p* = .718; Table S24). Finally, as in the main analysis, there was no difference in prediction performance of personality traits with region-specific SC-FC coupling between trait-relevant and trait-irrelevant tasks (B-NMA; 5-fold internal cross-validation: Table S14; cross-sample model generalization test: Table S22). For further validation results see Tables S13 – S24.

### 3.5. Post-hoc analyses

#### 3.5.1. Task performance was not associated with personality traits

To test whether the implemented tasks led to personality trait-dependent variations in behavior, individual performance scores on all tasks but the motor task were computed according to Greene et al. (2020) with available data (*N* = 737) and set in relation (Pearson correlation) to personality trait scores. No significant associations were observed (Table S1; Bonferroni corrected threshold for 66 comparisons: *p* < .00076).

#### 3.5.2. SC-FC coupling better predicts intelligence than personality traits

Recently, it was demonstrated that individual general intelligence scores, computed as latent *g*-factor from 12 cognitive measures (Table S25) could be significantly predicted from resting-state, task-induced and task-combined SC-FC coupling in the same sample (Table S26; Popp et al., 2024, 2025). We compared the absolute difference in prediction performance between two models (resting-state/task-specific B-NMA or E-NMA) trained on observed intelligence versus personality trait scores (Δl*r_observed_*l) to the absolute difference in prediction performance obtained when the same models were trained on permuted scores (Δl*r_permuted_*l). Importantly, Δl*r_permuted_*l was computed 1000 times, to determine how often it exceeded Δl*r_observed_*l (significant *p* < .05). For all tasks but the motor task, prediction of general intelligence was descriptively better than of any personality trait, with some comparisons reaching significance (Figure 2E; Table S27).

## 4. Discussion

### 4.1. Summary

This study investigates the relationship between individual differences in the Big Five personality traits and SC-FC coupling during resting state and tasks, differing in their trait-relevance. We used data from the Human Connectome Project (*N_main_* = 532; *N_lockbox_* = 232; Van Essen et al., 2013), personality traits were assessed with the NEO-FFI, and SC-FC coupling was operationalized with one similarity and three communication measures. During unconstrained resting state, personality traits were not associated with SC-FC coupling. However, constraining the situational context led to significant associations: Brain-average coupling during the social cognition task was positively correlated with agreeableness, and conscientiousness could be significantly predicted from task-combined region-specific SC-FC coupling.

### 4.2. Personality traits are not associated with SC-FC coupling during resting state

We observed no significant relationship between personality traits and SC-FC coupling during resting state. This was unexpected, as resting-state SC-FC coupling integrates SC with resting-state FC and both have independently been linked to personality traits (Dubois et al., 2018; Hsu et al., 2018; Nostro et al., 2018; Ueda et al., 2018). Those previously observed associations seem plausible, as due to the temporal stability of personality traits, their neural correlates are assumed to persist in the absence of stimulation (Markett et al., 2018). A potential explanation for this unexpected observation could be the reduced reliability of our composite measure, as it is limited by the multiplication of both modalities’ measurement errors. Although SC-FC coupling aims at offering mechanistic insight, its reduced reliability might hinder detection of brain-trait relationships – especially during resting state, where unconstrained, rapidly shifting self-generated thought may destabilize SC-FC interactions and obscure subtle neural patterns linked to personality traits.

### 4.3. Constraining situational context with tasks reveals significant associations between personality traits and SC-FC coupling

First, we observed a positive association between agreeableness and SC-FC coupling during a task involving social demands – specifically when SC-FC coupling was operationalized with search information, a measure reflecting the ease for diffusive signal flow between two regions (Seguin et al., 2020). This might suggest that more agreeable individuals process social cues more efficiently, supporting social adaptation and positive interpersonal relationships. However, this finding was not observed in the lockbox sample, thus requiring further investigation.

Second, conscientiousness could be significantly predicted when combining SC-FC coupling across tasks and considering brain region-specific differences in neural communication. All tasks involved some kind of instruction, requiring participants to follow rules. Given that conscientious individuals typically show strong self-discipline and top-down control over behavior and motivation (Roberts et al., 2014), they may excel at these task-general demands.

Our finding suggests that this could be reflected in region-specific adaptions of SC-FC coupling. Again, as those results only appeared in the main sample, this interpretation also remains tentative.

Generally, these findings suggest that trait-relevant tasks have the potential to unmask individual differences in SC-FC coupling associated with variations in specific personality traits. This extends prior research on intelligence, showing improved prediction under increased cognitive demand from both FC (Greene et al., 2018; Thiele et al., 2024) and SC-FC coupling (Popp et al., 2025). Further, it advances initial work in the personality domain, where brain-trait associations were influenced by situational context, without considering trait-relevance (Hardikar et al., 2024). Although most of our results could only be detected in the main sample and thus warrant cautious interpretation, our findings contribute to the literature twofold: First, they indicate that well-selected tasks can amplify individual differences in trait-related neural characteristics by engaging trait-relevant neural circuitry, thereby reinforcing recent recommendations to optimize neuroimaging effect sizes by aligning tasks with constructs of interest (DeYoung et al., 2025). Second, they extend personality theories emphasizing the critical impact of situational context for detecting personality-related behavioral differences, such as the Trait Activation Theory (Tett & Burnett, 2003; Tett & Guterman, 2000) and Cybernetic Big Five Theory (DeYoung, 2015), to the neural level.

### 4.4. No consistent effect of trait-relevance

Beyond findings outlined above, our analysis revealed no overall effects of trait-relevance on associations between personality traits and SC-FC coupling. While this could suggest a weaker impact of trait-relevance than expected, we consider it more plausible that fMRI tasks were not well-suited to engage these traits. Relying on open data, task trait-relevance could only be assessed post hoc - a clear limitation. Future studies should explicitly design tasks to engage the trait of interest and naturalistic data like movie scenes should be considered, as their rich sensory and narrative content engages cognitive, social, and emotional processes resembling real-life experiences (Sonkusare et al., 2019).

### 4.5. Neural effects of trait-relevance without effects in behavior: Proposal of a threshold theory

Associations between personality traits and neural characteristics during trait-relevant tasks were not reflected in task performance. If this is confirmed in future research, we propose that the exposure to trait-relevant contexts initially induces adaptive changes in brain connectivity, optimizing further processing of trait-relevant cues, and that behavioral manifestations only occur when a certain threshold is passed. The position of this threshold, as well as the nature of situational cues and resulting behavior, likely depends on a person’s trait level. Although preliminary, our findings encourage further testing of this Brain-Personality Threshold Theory (BPTT).

### 4.6. Intelligence can be better predicted from SC-FC coupling than personality traits

Our post-hoc analysis revealed higher prediction performance for individual intelligence than for personality traits. This aligns with the higher reliability of intelligence tests compared to self-report personality assessments (Colom, 2004; Viswesvaran & Ones, 2000), and highlights the need for reliable performance-based personality measures with reduced subjectivity (e.g., Implicit Association Test; Grumm & Collani, 2007). Alternatively, it could imply a greater role of SC-FC coupling in intelligence, while the neural implementation of personality traits may be more variable and harder to capture with the available tasks.

## 5. Limitations and future directions

Beyond the previously discussed limitations, i.e., restricted measure reliability and vague trait-relevance of tasks, our data prevented analyses of SC-FC coupling in subcortical structures (despite their known implication in personality; DeYoung, 2010) and similar results were largely absent in the lockbox sample. As a small sample size could be an explanation for the latter, replication in larger, independent cohorts is required. More broadly, this study faced common challenges in personality neuroscience, including the assumption that the psychometrically defined, multi-faceted Big Five personality traits can be directly mapped onto similar neurobiological mechanisms (Yarkoni, 2015).

## 6. Conclusion

Our study reveals that selecting trait-relevant tasks can foster detecting neural substrates of personality traits. We propose the Brain-Personality Threshold Theory (BPTT) to explain brain-personality associations in the absence of behavioral relationships: Trait-relevant contexts initially amplify trait-related differences in neural circuitry, which only manifest behaviorally if a certain threshold is exceeded.

## Author contributions

**Johanna L. Popp:** Conceptualization, Data Curation, Formal analysis, Funding acquisition, Methodology, Visualization, Writing – original draft**. Jonas A. Thiele:** Methodology, Writing – review and editing. **Joshua Faskowitz:** Data Curation, Methodology, Resources, Writing – review and editing. **Caio Seguin:** Methodology, Writing – review and editing. **Olaf Sporns:** Methodology, Writing – review and editing. **Kirsten Hilger:** Conceptualization, Funding acquisition, Methodology, Resources, Supervision, Writing – original draft.

## Acknowledgements

We thank the Human Connectome Project, WU-Minn Consortium (Principal Investigators: David Van Essen and Kamil Ugurbil; 1U554MH091657), funded by the 16 NIH Institutes and Centers supporting the NIH Blueprint for Neuroscience Research and by the McDonnell Center for Systems Neuroscience at Washington University for providing data.

## Ethics statement

The Washington University Institutional Review Board (Van Essen et al., 2013) approved all procedures of the Human Connectome Project and all participants provided informed written consent in accordance with the principles of the declaration of Helsinki.

## Supplemental Material

Supplemental material is available in the online version of this article.

