## Supplementary Material for "Trait-Relevant Tasks Improve Personality Prediction from Structural-Functional Brain Network Coupling"

Short Title: Trait-Relevance Improves Personality Prediction

Johanna L. Popp[1]*, Jonas A. Thiele[1], Joshua Faskowitz[2], Caio Seguin[2,3], Olaf Sporns[2], Kirsten Hilger[1,4]*

[1] Department of Psychology I, Würzburg University, Marcusstr. 9-11, Würzburg D-97070, Germany

[2] Department of Psychological and Brain Sciences, Indiana University, 1101 E. 10th St., Bloomington, IN 47405-7007, USA

[3] Department of Psychiatry, The University of Melbourne, Grattan Street, Parkville, Victoria 3010, Australia

[4] Department of Psychology, Vinzenz Pallotti University, Pallottistr. 3, Vallendar D-56179, Germany

*** Correspondence**

Johanna L. Popp, Department of Psychology I, Würzburg University, Marcusstr. 9-11, Würzburg D-97070, Germany. **and**
Kirsten Hilger, Department of Psychology, Vinzenz Pallotti University, Pallottistr. 3, Vallendar D-56179, Germany.

**Please note that some figures and paragraphs in this Supplemental Material have previously been published in related work (Popp et al., 2025), as the same sample and similar analysis pipelines were used. These parts are explicitly labeled.**

**Table S1**

Associations between in-scanner task performance and personality scores

| Task performance | Task performance *r* (*p*) EMO | GAM | LAN | REL | SOC | WM |
|---|---|---|---|---|---|---|
| **GAM** | .08 (.025) | | | | | |
| **LAN** | .10 (.010) | -.04 (.244) | - | | | |
| **REL** | **.15 (< .001)** | **-.13 (< .001)** | **.37 (< .001)** | - | | |
| **SOC** | .04 (.266) | -.05 (.170) | .07 (.057) | **.20 (< .001)** | - | |
| **WM** | .12 (.001) | -.12 (.002) | **.34 (< .001)** | **.48 (< .001)** | **.21 (<.001)** | - |

| Trait | Task performance *r* (*p*) EMO | GAM | LAN | REL | SOC | WM |
|---|---|---|---|---|---|---|
| **A** | .05 (.142) | .00 (.989) | .00 (.978) | .03 (.379) | .00 (.961) | .00 (.975) |
| **O** | -.03 (.436) | .01 (.893) | .08 (.034) | .11 (.002) | -.03 (.399) | .11 (.002) |
| **C** | -.01 (.720) | -.03 (.436) | -.06 (.125) | -.07 (.053) | .01 (.833) | -.06 (.089) |
| **N** | -.05 (.179) | .09 (.017) | -.07 (.073) | -.04 (.328) | -.05 (.147) | -.11 (.003) |
| **E** | -.05 (.193) | -.05 (.165) | -.05 (.217) | -.01 (.720) | -.02 (.657) | .07 (.061) |
| ***g*-factor** | **.13 (< .001)** | -.07 (.07) | **.45 (< .001)** | **.45 (<.001)** | **.23 (< .001)** | **.49 (<.001)** |

| Trait | Trait *r* (*p*) A | O | C | N | E |
|---|---|---|---|---|---|
| **A** | - | | | | |
| **O** | .05 (.140) | - | | | |
| **C** | **.19 (<.001)** | **-.17 (<.001)** | - | | |
| **N** | **-.26 (<.001)** | .04 (.270) | **-.42 (<.001)** | - | |
| **E** | **.27 (<.001)** | .09 (.013) | **.25 (<.001)** | **-.36 (<.001)** | - |
| ***g*-factor** | .07 (.044) | **.29 (<.001)** | -.11 (.002) | -.07 (.071) | -.05 (.150) |

*Note:* Pearson correlations (*r*) between personality, intelligence and available task performance scores for all participants with available data ($N$ = 737). The performance scores were computed as outlined in Greene et al. (2020) based on measures provided by the Human Connectome Project. No performance scores were available for the motor task. Significant associations when corrected for multiple comparisons (Bonferroni corrected threshold for 66 unique correlations: $p < .00076$) are presented in bold. Preregistered trait-relevant tasks in the bottom left section are highlighted in dark grey. EMO = Emotion processing task; GAM = Gambling task; LAN = Language task; REL = Relational processing task; SOC = Social cognition task; WM = Working memory task; A = Agreeableness; O = Openness to experience; C = Conscientiousness; N = Neuroticism; E = Extraversion.

**Table S2**

Relationship between personality scores and brain-average SC-FC coupling during resting state

| Resting state | | | | | |
|---|---|---|---|---|---|
| **SC-FC coupling measure** | **Personality trait *r* (*p*)** | | | | |
| | **A** | **O** | **C** | **N** | **E** |
| **PL** | -.02 (.689) | -.04 (.323) | .01 (.848) | .01 (.742) | .03 (.438) |
| **G** | -.02 (.598) | -.05 (.256) | -.02 (.579) | .03 (.428) | .02 (.637) |
| **CoS** | -.01 (.898) | -.04 (.337) | -.01 (.749) | .03 (.519) | .04 (.371) |
| **SI** | .01 (.773) | .00 (.921) | -.04 (.400) | .02 (.668) | .02 (.699) |

*Note*: Partial correlations between personality scores and brain-average coupling values (measure-specific averages of coupling values from all brain regions) during resting state controlled for age, gender, handedness, and in-scanner head motion. No associations passed the Bonferroni-corrected threshold (for four comparisons; $p < .0125$). PL = Path length; G = Communicability; CoS = Cosine similarity; SI = Search information; A = Agreeableness; O = Openness to experience; C = Conscientiousness; N = Neuroticism; E = Extraversion.

**Table S3**

Performance when predicting personality scores from brain region-specific SC-FC coupling

| **Prediction model performance** *r* (*p*) | **Personality trait** | | | | |
|---|---|---|---|---|---|
| **Condition** | **A** | **O** | **C** | **N** | **E** |
| **RES** | -.01 (.617) | -.03 (.725) | .07 (.072) | -.01 (.637) | .06 (.109) |
| **EMO** | .01 (.405) | .03 (.236) | .03 (.289) | -.06 (.924) | .00 (.493) |
| **GAM** | -.04 (.850) | .05 (.137) | **.10 (.032)*** | -.03 (.723) | .05 (.128) |
| **LAN** | .03 (.279) | .08 (.071) | **.07 (.048)*** | .06 (.085) | .04 (.137) |
| **MOT** | -.01 (.625) | .02 (.357) | .08 (.053) | -.03 (.737) | .03 (.215) |
| **REL** | .04 (.170) | -.08 (.958) | **.08 (.045)*** | .01 (.397) | .03 (.288) |
| **SOC** | **.08 (.042)*** | .07 (.105) | .07 (.056) | -.06 (.932) | **.08 (.028)*** |
| **WM** | .06 (.086) | .05 (.128) | **.09 (.044)*** | .06 (.090) | .03 (.238) |

*Note:* Personality trait predictions from brain region-specific coupling information during single tasks were based on the Basic NMA Model. Significance of the prediction performance was assessed with non-parametric permutation tests. *P*-values indicating significant associations uncorrected for multiple comparisons are marked with one asterisk (* = $p < .05$). Trait-relevant prediction models are highlighted in grey. RES = Resting state; EMO = Emotion processing task; GAM = Gambling task; LAN = Language task; MOT = Motor task; REL = Relational processing task; SOC = Social cognition task; WM = Working memory task; A = Agreeableness; O = Openness to experience; C = Conscientiousness; N = Neuroticism; E = Extraversion.

**Table S4**

Relationship between personality scores and brain-average SC-FC coupling during the emotion processing task

| Emotion processing task | | | | | |
|---|---|---|---|---|---|
| **SC-FC coupling measure** | **Personality trait *r* (*p*)** | | | | |
| | **A** | **O** | **C** | **N** | **E** |
| **PL** | -.01 (.882) | .03 (.525) | -.05 (.246) | .01 (.768) | -.04 (.402) |
| **G** | .00 (.913) | -.01 (.900) | -.10 (.018) | .01 (.888) | -.02 (.589) |
| **CoS** | .00 (.914) | .00 (.979) | -.09 (.034) | .01 (.781) | -.03 (.530) |
| **SI** | .04 (.360) | .05 (.241) | -.06 (.170) | .01 (.899) | -.06 (.139) |

*Note*: Partial correlations between personality scores and brain-average coupling values (measure-specific averages of coupling values from all brain regions) during the emotion processing task controlled for age, gender, handedness, and in-scanner head motion. No associations passed the Bonferroni-corrected threshold (for four comparisons; $p < .0125$). PL = Path length; G = Communicability; CoS = Cosine similarity; SI = Search information; A = Agreeableness; O = Openness to experience; C = Conscientiousness; N = Neuroticism; E = Extraversion.

**Table S5**

Relationship between personality scores and brain-average SC-FC coupling during the gambling task

| Gambling task | | | | | |
|---|---|---|---|---|---|
| **SC-FC coupling measure** | **Personality trait *r* (*p*)** | | | | |
| | **A** | **O** | **C** | **N** | **E** |
| **PL** | .02 (.687) | -.05 (.219) | .01 (.818) | .04 (.322) | -.03 (.468) |
| **G** | .01 (.870) | -.08 (.062) | -.04 (.329) | .07 (.133) | -.04 (.309) |
| **CoS** | .02 (.704) | -.07 (.104) | -.02 (.618) | .06 (.192) | -.04 (.330) |
| **SI** | .06 (.169) | -.01 (.882) | -.01 (.770) | .05 (.212) | -.07 (.093) |

*Note*: Partial correlations between personality scores and brain-average coupling values (measure-specific averages of coupling values from all brain regions) during the gambling task controlled for age, gender, handedness, and in-scanner head motion. No associations passed the Bonferroni-corrected threshold (for four comparisons; $p < .0125$). PL = Path length; G = Communicability; CoS = Cosine similarity; SI = Search information; A = Agreeableness; O = Openness to experience; C = Conscientiousness; N = Neuroticism; E = Extraversion.

**Table S6**

Relationship between personality scores and brain-average SC-FC coupling during the language task

| Language task | | | | | |
|---|---|---|---|---|---|
| **SC-FC coupling measure** | **Personality trait *r* (*p*)** | | | | |
| | **A** | **O** | **C** | **N** | **E** |
| **PL** | .02 (.569) | -.08 (.061) | -.00 (.965) | .05 (.253) | -.00 (.929) |
| **G** | .03 (.461) | -.10 (.025) | -.01 (.780) | .04 (.353) | .02 (.692) |
| **CoS** | .03 (.469) | -.10 (.016) | .01 (.863) | .03 (.511) | .02 (.583) |
| **SI** | .05 (.291) | -.06 (.141) | -.01 (.831) | .05 (.263) | -.04 (.408) |

*Note*: Partial correlations between personality scores and brain-average coupling values (measure-specific averages of coupling values from all brain regions) during the language task controlled for age, gender, handedness, and in-scanner head motion. No associations passed the Bonferroni-corrected threshold (for four comparisons; $p < .0125$). PL = Path length; G = Communicability; CoS = Cosine similarity; SI = Search information; A = Agreeableness; O = Openness to experience; C = Conscientiousness; N = Neuroticism; E = Extraversion.

**Table S7**

Relationship between personality scores and brain-average SC-FC coupling during the motor task

| Motor task | | | | | |
|---|---|---|---|---|---|
| **SC-FC coupling measure** | **Personality trait *r* (*p*)** | | | | |
| | **A** | **O** | **C** | **N** | **E** |
| **PL** | .04 (.318) | -.06 (.148) | -.04 (.328) | .02 (.614) | .02 (.674) |
| **G** | .06 (.176) | -.07 (.108) | -.06 (.144) | .01 (.786) | .03 (.548) |
| **CoS** | .06 (.139) | -.07 (.124) | -.06 (.147) | .02 (.724) | .03 (.512) |
| **SI** | .08 (.055) | -.03 (.442) | -.08 (.078) | .03 (.491) | -.01 (.899) |

*Note*: Partial correlations between personality scores and brain-average coupling values (measure-specific averages of coupling values from all brain regions) during the motor task controlled for age, gender, handedness, and in-scanner head motion. No associations passed the Bonferroni-corrected threshold (for four comparisons; $p < .0125$). PL = Path length; G = Communicability; CoS = Cosine similarity; SI = Search information; A = Agreeableness; O = Openness to experience; C = Conscientiousness; N = Neuroticism; E = Extraversion.

**Table S8**

Relationship between personality scores and brain-average SC-FC coupling during the relational processing task

| Relational processing task | | | | | |
|---|---|---|---|---|---|
| **SC-FC coupling measure** | **Personality trait *r* (*p*)** | | | | |
| | **A** | **O** | **C** | **N** | **E** |
| **PL** | -.00 (.966) | .02 (.590) | -.06 (.199) | .05 (.250) | -.08 (.076) |
| **G** | .01 (.880) | -.00 (.980) | -.10 (.025) | .06 (.200) | -.08 (.082) |
| **CoS** | .02 (.664) | .00 (.932) | -.08 (.075) | .05 (.300) | -.06 (.149) |
| **SI** | .03 (.456) | .04 (.314) | -.05 (.296) | .03 (.475) | -.07 (.110) |

*Note*: Partial correlations between personality scores and brain-average coupling values (measure-specific averages of coupling values from all brain regions) during the relational processing task controlled for age, gender, handedness, and in-scanner head motion. No associations passed the Bonferroni-corrected threshold (for four comparisons; $p < .0125$). PL = Path length; G = Communicability; CoS = Cosine similarity; SI = Search information; A = Agreeableness; O = Openness to experience; C = Conscientiousness; N = Neuroticism; E = Extraversion.

**Table S9**

Relationship between personality scores and brain-average SC-FC coupling during the social cognition task

| Social cognition task | | | | | |
|---|---|---|---|---|---|
| **SC-FC coupling measure** | **Personality trait *r* (*p*)** | | | | |
| | **A** | **O** | **C** | **N** | **E** |
| **PL** | .09 (.047) | -.06 (.175) | .00 (.956) | .03 (.536) | .00 (.928) |
| **G** | .05 (.260) | -.09 (.039) | -.04 (.321) | .03 (.441) | -.03 (.513) |
| **CoS** | .07 (.112) | -.09 (.046) | -.03 (.531) | .04 (.388) | -.02 (.700) |
| **SI** | **.12 (.006)*** | -.04 (.364) | .02 (.585) | .02 (.673) | -.03 (.560) |

*Note*: Partial correlations between personality scores and brain-average coupling values (measure-specific averages of coupling values from all brain regions) during the social cognition task controlled for age, gender, handedness, and in-scanner head motion. Significant associations passing the Bonferroni-corrected threshold (for four comparisons) are marked with an asterisk (* = $p$ < .0125). PL = Path length; G = Communicability; CoS = Cosine similarity; SI = Search information; A = Agreeableness; O = Openness to experience; C = Conscientiousness; N = Neuroticism; E = Extraversion.

**Table S10**

Relationship between personality scores and brain-average SC-FC coupling during the working memory task

| Working memory task | | | | | |
|---|---|---|---|---|---|
| SC-FC coupling measure | Personality trait *r* (*p*) | | | | |
| | A | O | C | N | E |
| **PL** | .02 (.676) | -.05 (.210) | -.07 (.114) | .02 (.681) | -.06 (.196) |
| **G** | .03 (.447) | -.10 (.027) | -.10 (.023) | .04 (.333) | -.07 (.113) |
| **CoS** | .03 (.442) | -.08 (.075) | -.10 (.025) | .04 (.317) | -.06 (.205) |
| **SI** | .07 (.133) | -.02 (.616) | -.09 (.045) | .02 (.631) | -.06 (.161) |

*Note*: Partial correlations between personality scores and brain-average coupling values (measure-specific averages of coupling values from all brain regions) during the working memory task controlled for age, gender, handedness, and in-scanner head motion. No associations passed the Bonferroni-corrected threshold (for four comparisons; $p < .0125$). PL = Path Length; G = Communicability; CoS = Cosine similarity; SI = Search information; A = Agreeableness; O = Openness to experience; C = Conscientiousness; N = Neuroticism; E = Extraversion.

**Table S11**

Performance when predicting personality traits from brain region-specific SC-FC coupling

| Personality trait | Prediction performance *r* (*p*) | |
|---|---|---|
| | All task conditions | Resting state |
| **A** | .04 (.187) | -.01 (.617) |
| **O** | .06 (.085) | -.03 (.725) |
| **C** | **.14 (.001)**** | .07 (.072) |
| **N** | .02 (.346) | -.01 (.637) |
| **E** | .06 (.051) | .06 (.109) |

*Note:* Personality trait prediction from task-combined region-specific coupling information was performed with the Expanded NMA Model, while personality trait prediction from brain region-specific coupling information during the resting state was based on the Basic NMA Model (see Methods). Significance of the prediction performance was assessed with non-parametric permutation tests. *P*-values indicating significant associations uncorrected for multiple comparisons are marked with one asterisk (* = $p < .05$), while significant associations passing the Bonferroni-corrected threshold (for eleven comparisons) are marked with two asterisks (** = $p < .005$). A = Agreeableness; O = Openness to experience; C = Conscientiousness; N = Neuroticism; E = Extraversion.

**Table S12**

Comparison of the performance when predicting personality traits from brain region-specific SC-FC coupling combined from trait-relevant vs. trait-irrelevant task conditions

| Personality trait | Prediction performance *r* (*p*) | |
|---|---|---|
| | **Trait-relevant tasks** | **Trait-irrelevant tasks** |
| **A** | .06 (.098) | .02 (.336) |
| **O** | .06 (.088) | .07 (.068) |
| **C** | **.10 (.032)*** | .08 (.066) |
| **N** | -.03 (.723) | -.00 (545) |
| **E** | **.09 (.046)*** | .02 (.294) |

*Note:* Personality trait prediction from combined region-specific coupling information across all trait-relevant and across all trait-irrelevant task conditions were performed with the Latent NMA Model (see Methods). Significance of the prediction performance was assessed with non-parametric permutation tests. *P*-values indicating significant associations uncorrected for multiple comparisons are marked with one asterisk (* = $p < .05$). No associations passed the Bonferroni-corrected threshold (for eleven comparisons; $p < .005$). Please note that there was only one trait-relevant task for conscientiousness and neuroticism, which is why in these cases the Basic NMA Model was applied. A = Agreeableness; O = Openness to experience; C = Conscientiousness; N = Neuroticism; E = Extraversion.

**Table S13**

Relationship between personality scores and brain-average SC-FC coupling during resting state in the lockbox sample

| **Resting state** | | | | | |
|---|---|---|---|---|---|
| **SC-FC coupling measure** | **Personality trait *r* (*p*)** | | | | |
| | **A** | **O** | **C** | **N** | **E** |
| **PL** | .02 (.818) | .16 (.015) | -.10 (.129) | -.05 (.414) | .07 (.307) |
| **G** | .00 (.997) | .16 (.017) | -.07 (.285) | -.05 (.491) | .04 (.516) |
| **CoS** | .02 (.728) | .15 (.028) | -.10 (.147) | -.05 (.455) | .04 (.582) |
| **SI** | -.04 (.577) | **.17 (.009)*** | -.13 (.047) | -.06 (.357) | .07 (.296) |

*Note*: Lockbox Sample $N = 232$. Partial correlations between personality scores and brain-average coupling values (measure-specific averages of coupling values from all brain regions) during resting state controlled for age, gender, handedness, and in-scanner head motion. Significant associations passing the Bonferroni-corrected threshold (for four comparisons) are marked with an asterisk (* = $p < .0125$). PL = Path length; G = Communicability; CoS = Cosine similarity; SI = Search information; A = Agreeableness; O = Openness to experience; C = Conscientiousness; N = Neuroticism; E = Extraversion.

**Table S14**

Performance when predicting personality scores from brain region-specific SC-FC coupling in the lockbox sample

| **Prediction model performance** *r* (*p*) | **Personality trait** | | | | |
|---|---|---|---|---|---|
| **Condition** | **A** | **O** | **C** | **N** | **E** |
| **RES** | .00 (.522) | .06 (.225) | -.02 (.623) | -.10 (.923) | .08 (.116) |
| **EMO** | -.03 (.648) | .10 (.116) | .08 (.157) | .02 (.405) | .04 (.285) |
| **GAM** | .02 (.367) | -.01 (.556) | .00 (.533) | -.19 (.990) | .00 (.543) |
| **LAN** | -.02 (.583) | **.13 (.036)*** | -.03 (.657) | -.04 (.695) | .08 (.131) |
| **MOT** | .05 (.251) | .03 (.275) | -.04 (.738) | -.10 (.906) | -.02 (.638) |
| **REL** | .00 (.481) | .03 (.333) | .05 (.198) | -.10 (.936) | .11 (.071) |
| **SOC** | -.04 (.706) | .05 (.206) | .02 (.410) | -.08 (.873) | .00 (.503) |
| **WM** | .10 (.116) | **.16 (.016)*** | **.16 (.022)*** | .08 (.087) | -.04 (.745) |

*Note:* Personality trait predictions from brain region-specific coupling information during single tasks were based on the Basic NMA Model. Significance of the prediction performance was assessed with non-parametric permutation tests. *P*-values indicating significant associations uncorrected for multiple comparisons are marked with one asterisk (* = $p < .05$). Trait-relevant prediction models are highlighted in grey. RES = Resting state; EMO = Emotion processing task; GAM = Gambling task; LAN = Language task; MOT = Motor task; REL = Relational processing task; SOC = Social cognition task; WM = Working memory task; A = Agreeableness; O = Openness to experience; C = Conscientiousness; N = Neuroticism; E = Extraversion.

**Table S15**

Relationship between personality scores and brain-average SC-FC coupling during the emotion processing task in the lockbox sample

| Emotion processing task | | | | | |
|---|---|---|---|---|---|
| **SC-FC coupling measure** | **Personality trait *r* (*p*)** | | | | |
| | **A** | **O** | **C** | **N** | **E** |
| **PL** | .00 (.968) | .09 (.196) | -.01 (.877) | .03 (.607) | .03 (.654) |
| **G** | .01 (.846) | .08 (.243) | .03 (.676) | .05 (.418) | .02 (.771) |
| **CoS** | .00 (.967) | .06 (.329) | .00 (.957) | .05 (.413) | .03 (.634) |
| **SI** | -.05 (.478) | .09 (.171) | -.05 (.477) | .02 (.744) | .02 (.768) |

*Note*: Lockbox sample $N$ = 232. Partial correlations between personality scores and brain-average coupling values (measure-specific averages of coupling values from all brain regions) during the emotion processing task controlled for age, gender, handedness, and in-scanner head motion. No associations passed the Bonferroni-corrected threshold (for four comparisons; $p < .0125$). PL = Path length; G = Communicability; CoS = Cosine similarity; SI = Search information; A = Agreeableness; O = Openness to experience; C = Conscientiousness; N = Neuroticism; E = Extraversion.

**Table S16**

Relationship between personality scores and brain-average SC-FC coupling during the gambling task in the lockbox sample

| Gambling task | | | | | |
|---|---|---|---|---|---|
| **SC-FC coupling measure** | **Personality trait *r* (*p*)** | | | | |
| | **A** | **O** | **C** | **N** | **E** |
| **PL** | -.06 (.382) | .04 (.589) | -.05 (.464) | .07 (.280) | .08 (.214) |
| **G** | -.04 (.591) | .03 (.672) | -.02 (.752) | .10 (.115) | .06 (.337) |
| **CoS** | -.04 (.585) | .03 (.619) | -.04 (.504) | .10 (.133) | .07 (.313) |
| **SI** | -.03 (.651) | .05 (.429) | -.08 (.248) | .05 (.492) | .07 (.297) |

*Note*: Lockbox Sample $N = 232$. Partial correlations between personality scores and brain-average coupling values (measure-specific averages of coupling values from all brain regions) during the gambling task controlled for age, gender, handedness, and in-scanner head motion. No associations passed the Bonferroni-corrected threshold (for four comparisons; $p < .0125$). PL = Path Length; G = Communicability; CoS = Cosine similarity; SI = Search information; A = Agreeableness; O = Openness to experience; C = Conscientiousness; N = Neuroticism; E = Extraversion.

**Table S17**

Relationship between personality scores and brain-average SC-FC coupling during the language task in the lockbox sample

| Language task | | | | | |
|---|---|---|---|---|---|
| **SC-FC coupling measure** | **Personality traits *r* (*p*)** | | | | |
| | **A** | **O** | **C** | **N** | **E** |
| **PL** | .09 (.196) | -.05 (.469) | -.04 (.590) | .00 (.979) | .10 (.163) |
| **G** | .08 (.246) | -.04 (.539) | -.02 (.804) | .02 (.736) | .11 (.101) |
| **CoS** | .09 (.156) | -.05 (.420) | -.04 (.595) | .03 (.695) | .11 (.112) |
| **SI** | .06 (.382) | -.05 (.456) | -.06 (.408) | .01 (.845) | .08 (.228) |

*Note*: Lockbox Sample $N$ = 232. Partial correlations between personality scores and brain-average coupling values (measure-specific averages of coupling values from all brain regions) during the language task controlled for age, gender, handedness, and in-scanner head motion. No associations passed the Bonferroni-corrected threshold (for four comparisons; $p < .0125$). PL = Path length; G = Communicability; CoS = Cosine similarity; SI = Search information; A = Agreeableness; O = Openness to experience; C = Conscientiousness; N = Neuroticism; E = Extraversion.

**Table S18**

Relationship between personality scores and brain-average SC-FC coupling during the motor task in the lockbox sample

| Motor task | | | | | |
|---|---|---|---|---|---|
| **SC-FC coupling measure** | **Personality traits *r* (*p*)** | | | | |
| | **A** | **O** | **C** | **N** | **E** |
| **PL** | -.10 (.150) | .03 (.692) | -.10 (.147) | .10 (.121) | .01 (.842) |
| **G** | -.07 (.305) | .00 (.976) | -.03 (.636) | .10 (.132) | .02 (.779) |
| **CoS** | -.07 (.262) | .01 (.824) | -.07 (.315) | .12 (.076) | .00 (.985) |
| **SI** | -.09 (.158) | .05 (.420) | -.10 (.139) | .10 (.129) | .04 (.594) |

*Note*: Lockbox Sample $N$ = 232. Partial correlations between personality scores and brain-average coupling values (measure-specific averages of coupling values from all brain regions) during the motor task controlled for age, gender, handedness, and in-scanner head motion. No associations passed the Bonferroni-corrected threshold (for four comparisons; $p < .0125$). PL = Path length; G = Communicability; CoS = Cosine similarity; SI = Search information; A = Agreeableness; O = Openness to experience; C = Conscientiousness; N = Neuroticism; E = Extraversion.

**Table S19**

Relationship between personality scores and brain-average SC-FC coupling during the relational processing task in the lockbox sample

| Relational processing task | | | | | |
|---|---|---|---|---|---|
| **SC-FC coupling measure** | **Personality trait *r* (*p*)** | | | | |
| | **A** | **O** | **C** | **N** | **E** |
| **PL** | .01 (.906) | .06 (.364) | .02 (.715) | .04 (.507) | .07 (.295) |
| **G** | .02 (.741) | .05 (.497) | .06 (.394) | .06 (.392) | .02 (.746) |
| **CoS** | .03 (.624) | .05 (.454) | .03 (.608) | .06 (.364) | .02 (.738) |
| **SI** | .00 (.942) | .06 (.331) | -.02 (.714) | .04 (.536) | .08 (.252) |

*Note*: Lockbox Sample $N$ = 232. Partial correlations between personality scores and brain-average coupling values (measure-specific averages of coupling values from all brain regions) during the relational processing task controlled for age, gender, handedness, and in-scanner head motion. No associations passed the Bonferroni-corrected threshold (for four comparisons; $p < .0125$). PL = Path length; G = Communicability; CoS = Cosine similarity; SI = Search information; A = Agreeableness; O = Openness to experience; C = Conscientiousness; N = Neuroticism; E = Extraversion.

**Table S20**

Relationship between personality scores and brain-average SC-FC coupling during the social cognition task in the lockbox sample (HCP232)

| Social cognition task | | | | | |
|---|---|---|---|---|---|
| **SC-FC coupling measure** | **Personality trait *r* (*p*)** | | | | |
| | **A** | **O** | **C** | **N** | **E** |
| **PL** | .02 (.784) | .01 (.848) | -.12 (.065) | -.03 (.650) | .05 (.468) |
| **G** | .02 (.806) | .00 (.949) | -.11 (.102) | .02 (.815) | .01 (.842) |
| **CoS** | .02 (.713) | .01 (.868) | -.14 (.039) | .00 (.976) | .02 (.802) |
| **SI** | .00 (.956) | .03 (.609) | -.13 (.049) | -.05 (.428) | .03 (.616) |

*Note*: Lockbox Sample $N$ = 232. Partial correlations between personality scores and brain-average coupling values (measure-specific averages of coupling values from all brain regions) during the social cognition task controlled for age, gender, handedness, and in-scanner head motion. No associations passed the Bonferroni-corrected threshold (for four comparisons; $p$ < .0125). PL = Path length; G = Communicability; CoS = Cosine similarity; SI = Search information; A = Agreeableness; O = Openness to experience; C = Conscientiousness; N = Neuroticism; E = Extraversion.

**Table S21**

Relationship between personality scores and brain-average SC-FC coupling during the working memory task in the lockbox sample

| **Working memory task** | | | | | |
|---|---|---|---|---|---|
| **SC-FC coupling measure** | **Personality trait *r* (*p*)** | | | | |
| | **A** | **O** | **C** | **N** | **E** |
| **PL** | .08 (.259) | .05 (.494) | -.04 (.564) | .02 (.793) | .03 (.622) |
| **G** | .06 (.388) | -.02 (.775) | .04 (.591) | .06 (.364) | .02 (.745) |
| **CoS** | .08 (.208) | .01 (.906) | -.01 (.893) | .06 (.343) | .01 (.854) |
| **SI** | .08 (.223) | .12 (.073) | -.11 (.088) | .03 (.706) | .02 (.789) |

*Note*: Lockbox Sample $N = 232$. Partial correlations between personality scores and brain-average coupling values (measure-specific averages of coupling values from all brain regions) during the working memory task controlled for age, gender, handedness, and in-scanner head motion. No associations passed the Bonferroni-corrected threshold (for four comparisons; $p < .0125$). PL = Path length; G = Communicability; CoS = Cosine similarity; SI = Search information; A = Agreeableness; O = Openness to experience; C = Conscientiousness; N = Neuroticism; E = Extraversion.

**Table S22**

Performance when predicting personality scores from brain region-specific SC-FC coupling in the cross-sample model generalization test in the lockbox sample

| **Prediction model performance** *r* (*p*) | **Personality trait** | | | | |
|---|---|---|---|---|---|
| **Condition** | **A** | **O** | **C** | **N** | **E** |
| **RES** | -.03 (.892) | .06 (.170) | -.02 (.741) | **.05 (.017)*** | -.04 (.874) |
| **EMO** | -.05 (.992) | .04 (.146) | -.04 (.967) | **.09 (.003)**** | .00 (.508) |
| **GAM** | -.10 (.995) | -.03 (.860) | -.05 (.959) | -.04 (.824) | **.08 (.013)*** |
| **LAN** | -.01 (.626) | **.14 (< .001)**** | -.02 (.767) | .02 (.075) | -.02 (.702) |
| **MOT** | .02 (.291) | .04 (.114) | -.02 (.744) | **.09 (.020)*** | -.06 (.978) |
| **REL** | -.04 (.936) | **.06 (.030)*** | -.04 (.998) | -.06 (.983) | -.15 (.999) |
| **SOC** | -.01 (.723) | **.08 (.004)**** | .03 (.226) | .01 (.243) | .03 (.075) |
| **WM** | -.05 (.905) | **.18 (< .001)**** | .05 (.053) | .03 (.078) | -.05 (.966) |

*Note:* Personality trait predictions in the cross-sample model generalization test (HCP532 → HCP232) from brain region-specific coupling information during single tasks were based on the Basic NMA Model. Significance of the prediction performance was assessed with non-parametric permutation tests. *P*-values indicating significant associations uncorrected for multiple comparisons are marked with one asterisk (* = $p < .05$), while significant associations passing the Bonferroni-corrected threshold (for eleven comparisons) are marked with two asterisks (** = $p < .005$). RES = Resting state; EMO = Emotion processing task; GAM = Gambling task; LAN = Language task; MOT = Motor task REL = Relational processing task; SOC = Social cognition task; WM = Working memory task; ALL = All task conditions; A = Agreeableness; O = Openness to experience; C = Conscientiousness; N = Neuroticism; E = Extraversion.

**Table S23**

Comparison of the performance when predicting personality traits from brain region-specific SC-FC coupling combined from all seven task conditions vs. the resting state condition in the lockbox sample

| **Personality trait** | **Prediction model performance *r* (*p*)** | |
|---|---|---|
| | **All task conditions** | **Resting state** |
| **A** | **.13 (.024)*** | .00 (.522) |
| **O** | **.18 (.006)*** | .06 (.225) |
| **C** | .08 (.142) | -.02 (.623) |
| **N** | -.04 (.727) | -.10 (.923) |
| **E** | **.14 (.016)*** | .08 (.116) |

*Note:* Lockbox Sample *N* = 232. Personality trait predictions from combined region-specific coupling information across all task conditions were performed with the Expanded NMA Model, while personality trait predictions from brain region-specific coupling information during the resting-state condition were based on the Basic NMA Model (see Methods). Significance of the prediction performance was assessed with non-parametric permutation tests. *P*-values indicating significant associations uncorrected for multiple comparisons are marked with one asterisk (* = $p < .05$). No associations passed the Bonferroni-corrected threshold (for eleven comparisons; $p < .005$). A = Agreeableness; O = Openness to experience; C = Conscientiousness; N = Neuroticism; E = Extraversion.

**Table S24**

Comparison of the performance when predicting personality traits from brain region-specific SC-FC coupling combined from all seven task conditions vs. the resting-state condition in the cross-sample model generalization test in the lockbox sample

| Personality trait | Prediction model performance *r* (*p*) | |
|---|---|---|
| | **All task conditions** | **Resting state** |
| **A** | -.05 (.952) | -.03 (.892) |
| **O** | **.12 (<.001)**** | .06 (.170) |
| **C** | -.01 (.718) | -.02 (.741) |
| **N** | .04 (.105) | **.05 (.017)*** |
| **E** | -.08 (.994) | -.04 (.874) |

*Note:* Personality trait predictions in the cross-sample model generalization test (HCP532 → HCP232) from combined region-specific coupling information across all task conditions were performed with the Expanded NMA Model, while personality trait predictions from brain region-specific coupling information during the resting-state condition were based on the Basic NMA Model (see Methods). Significance of the prediction performance was assessed with non-parametric permutation tests. *P*-values indicating significant associations uncorrected for multiple comparisons are marked with one asterisk (* = $p < .05$), while significant associations passing the Bonferroni-corrected threshold (for eleven comparisons) are marked with two asterisks (** = $p < .005$). A = Agreeableness; O = Openness to experience; C = Conscientiousness; N = Neuroticism; E = Extraversion.

**Table S25**

Cognitive tests and measures used to calculate a latent *g*-factor as estimate of general intelligence

| Test | Instrument | Measure used as input for *g*-factor calculation |
|---|---|---|
| 1 | Episodic memory (Picture sequence memory) | PicSeq_Unadj |
| 2 | Executive function/cognitive flexibility (Dimensional change card sort) | CardSort_Unadj |
| 3 | Executive function/inhibition (Flanker task) | Flanker_Unadj |
| 4 | Fluid intelligence (Penn Progressive Matrices) | PMAT24_A_CR |
| 5 | Language/reading decoding (Oral reading recognition) | ReadEng_Unadj |
| 6 | Language/vocabulary comprehension (Picture vocabulary) | PicVocab_Unadj |
| 7 | Processing speed (Pattern completion processing speed) | ProcSpeed_Unadj |
| 8 | Self-regulation/impulsivity (Delay discounting) | DDisc_AUC_200 + DDisc_AUC_40K |
| 9 | Spatial orientation (Variable Short Penn Line Orientation Test) | VSPLOT_TC |
| 10 | Sustained attention (Short Penn Continuous Performance Test) | $\frac{SCPT\_TP + SCPT\_TN}{(SCPT\_TP + SCPT\_TN + SCPT\_FP + SCPT\_FN)SCPT\_TPRT}$ |
| 11 | Verbal episodic memory (Penn Word Memory Test) | IWRD_TOT |
| 12 | Working memory (List sorting) | ListSort_Unadj |

*Note:* Bi-factor analysis was performed to estimate a latent factor of general intelligence (Dubois et al., 2018; Schmid & Leiman, 1957). Analyses were conducted in a larger sample (*N* = 1186; Thiele et al., 2022) utilizing data from 12 cognitive measures (Barch et al., 2013). This table was adapted from Popp et al. (2025).

**Table S26**

Performance when predicting intelligence from brain region-specific SC-FC coupling

| FMRI condition | Prediction performance *r* (*p*) |
|---|---|
| **RES** | .18 (.002)** |
| **EMO** | .12 (.029)* |
| **GAM** | .18 (< .001)** |
| **LAN** | .15 (.001)** |
| **MOT** | .07 (.072) |
| **REL** | .12 (.014)* |
| **SOC** | .22 (< .001)** |
| **WM** | .25 (< .001)** |
| **ALL** | .27 (< .001)** |

*Note:* Intelligence was predicted with the Basic NMA Model based on brain region-specific coupling obtained during single tasks, while the Expanded NMA Model used task-combined region-specific coupling to predict intelligence (Popp et al., 2025). Significance of the prediction performance was assessed with non-parametric permutation tests. *P*-values indicating significant associations uncorrected for multiple comparisons are marked with one asterisk (* = $p < .05$), while significant associations passing the Bonferroni-corrected threshold (for nine comparisons) are marked with two asterisks (** = $p < .006$). RES = Resting state; EMO = Emotion processing task; GAM = Gambling task; LAN = Language task; MOT = Motor task; REL = Relational processing task; SOC = Social cognition task; WM = Working memory task; ALL = All task conditions.

**Table S27**

Assessment of significant differences in prediction performance between individual personality and general intelligence scores from brain region-specific SC-FC coupling

| | | Personality trait | | | | |
|---|---|---|---|---|---|---|
| | | A | O | C | N | E |
| B-NMA | RES | G > A<br>$\lvert r \rvert$ = .19<br>$p$ = .006* | G > O<br>$\lvert r \rvert$ = .20<br>$p$ = .004* | G > C<br>$\lvert r \rvert$ = .11<br>$p$ = .093 | G > N<br>$\lvert r \rvert$ = .19<br>$p$ = .006* | G > E<br>$\lvert r \rvert$ = .12<br>$p$ = .074 |
| | EMO | G > A<br>$\lvert r \rvert$ = .12<br>$p$ = .065 | G > O<br>$\lvert r \rvert$ = .09<br>$p$ = .100 | G > C<br>$\lvert r \rvert$ = .09<br>$p$ = .111 | G > N<br>$\lvert r \rvert$ = .18<br>$p$ = .007* | G > E<br>$\lvert r \rvert$ = .12<br>$p$ = .052 |
| | GAM | G > A<br>$\lvert r \rvert$ = .23<br>$p$ = .001* | G > O<br>$\lvert r \rvert$ = .14<br>$p$ = .036* | G > C<br>$\lvert r \rvert$ = .08<br>$p$ = .164 | G > N<br>$\lvert r \rvert$ = .21<br>$p$ = .003* | G > E<br>$\lvert r \rvert$ = .14<br>$p$ = .039* |
| | LAN | G > A<br>$\lvert r \rvert$ = .12<br>$p$ = .043* | G > O<br>$\lvert r \rvert$ = .07<br>$p$ = .208 | G > C<br>$\lvert r \rvert$ = .08<br>$p$ = .145 | G > N<br>$\lvert r \rvert$ = .08<br>$p$ = .144 | G > E<br>$\lvert r \rvert$ = .11<br>$p$ = .067 |
| | MOT | G > A<br>$\lvert r \rvert$ = .08<br>$p$ = .105 | G > O<br>$\lvert r \rvert$ = .05<br>$p$ = .208 | C > G<br>$\lvert r \rvert$ = .01<br>$p$ = .434 | G > N<br>$\lvert r \rvert$ = .09<br>$p$ = .072 | G > E<br>$\lvert r \rvert$ = .04<br>$p$ = .273 |
| | REL | G > A<br>$\lvert r \rvert$ = .08<br>$p$ = .120 | G > O<br>$\lvert r \rvert$ = .20<br>$p$ = .002* | G > C<br>$\lvert r \rvert$ = .04<br>$p$ = .299 | G > N<br>$\lvert r \rvert$ = .11<br>$p$ = .062 | G > E<br>$\lvert r \rvert$ = .10<br>$p$ = .092 |
| | SOC | G > A<br>$\lvert r \rvert$ = .14<br>$p$ = .069 | G > O<br>$\lvert r \rvert$ = .15<br>$p$ = .043* | G > C<br>$\lvert r \rvert$ = .15<br>$p$ = .051 | G > N<br>$\lvert r \rvert$ = .28<br>$p$ < .001* | G > E<br>$\lvert r \rvert$ = .13<br>$p$ = .063 |
| | WM | G > A<br>$\lvert r \rvert$ = .19<br>$p$ = .017* | G > O<br>$\lvert r \rvert$ = .20<br>$p$ = .013* | G > C<br>$\lvert r \rvert$ = .16<br>$p$ = .039* | G > N<br>$\lvert r \rvert$ = .19<br>$p$ = .016* | G > E<br>$\lvert r \rvert$ = .23<br>$p$ = .007* |
| E – NMA | ALL | G > A<br>$\lvert r \rvert$ = .23<br>$p$ = .001* | G > O<br>$\lvert r \rvert$ = .21<br>$p$ = .003* | G > C<br>$\lvert r \rvert$ = .13<br>$p$ = .038* | G > N<br>$\lvert r \rvert$ = .25<br>$p$ < .001* | G > E<br>$\lvert r \rvert$ = .20<br>$p$ = .003* |

*Note:* Significant differences in prediction performance between condition-specific models predicting general intelligence vs. each of the five personality traits from brain region-specific SC-FC coupling were assessed by comparing the absolute difference in prediction performance (correlation coefficients between predicted and observed intelligence scores) between two models trained on the observed intelligence scores versus trained on the personality trait scores ($\lvert r_{diffobserved} \rvert$; reported in table) to the absolute difference in prediction performance when the same two models were trained on the permuted scores ($\lvert r_{diffpermuted} \rvert$). *P*-values indicating significant model differences are marked with an asterisk (* = $p$ < .05;

|$r_{diffpermuted}$| was greater than |$r_{diffobserved}$| for less than 50/1.000 permutations). B-NMA = Basic NMA model; E-NMA = Expanded NMA model; RES = Resting state; EMO = Emotion processing task; GAM = Gambling task; LAN = Language task; MOT = Motor task; REL = Relational processing task; SOC = Social cognition task; WM = Working memory task; ALL = All task conditions; G = General intelligence; A = Agreeableness; O = Openness to experience; C = Conscientiousness; N = Neuroticism; E = Extraversion.

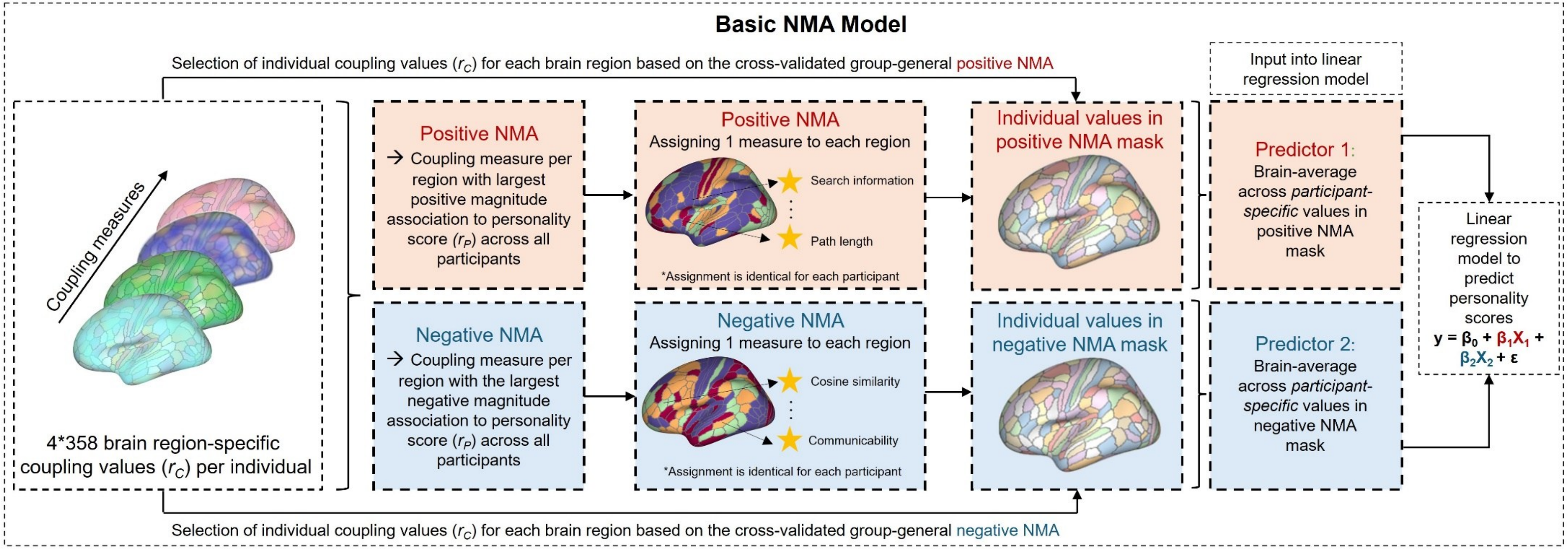


**Figure S1.** Visualization of the Basic NMA prediction model used predict individual personality scores from region-specific SC-FC coupling. Two global input features were generated by first determining a group-general positive and negative node-measure assignment mask (NMA) designating the measure with the largest positive magnitude association (positive NMA) and the largest negative magnitude association (negative NMA) to the respective personality score to a given brain region. These group-general masks were used to extract participant-specific regional coupling values ($r_c$) and ultimately, brain-averages across the participant-specific values in the positive NMA mask (predictor 1) and the negative NMA mask predictor 2) served as the two predictors for the 5-fold cross-validated multiple linear regression model. Importantly, NMAs built in the training samples were used to extract region-specific coupling values in the respective test samples to avoid data leakage between cross-validation folds. Prediction performance was assessed by correlating predicted and observed personality scores and significance was determined with non-parametric permutation tests. NMA = Node-measure assignment. This Figure was adapted from Popp et al. (2025).

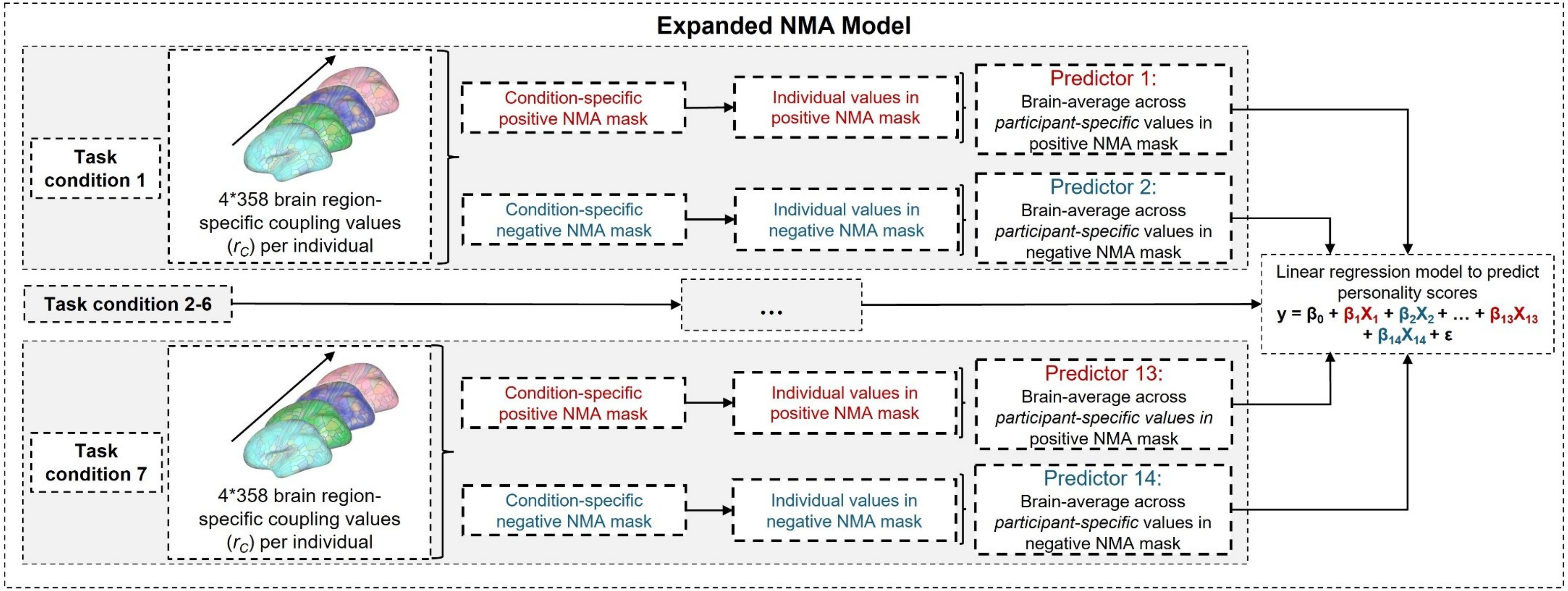


**Figure S2.** Visualization of the Expanded NMA prediction model used to predict individual personality scores from region-specific SC-FC coupling. For each task fMRI condition separately, group-general NMA masks were created based on the largest positive magnitude association (Positive NMA) and the largest negative magnitude association (Negative NMA) between region-specific coupling measures and the respective personality score. These group-general masks were used to extract participant-specific regional SC-FC coupling values ($r_C$) for each respective task-fMRI condition. Ultimately, brain-averages across the participant-specific values in the positive NMA mask (predictor 1) and the negative NMA mask (predictor 2) per task fMRI condition were used as model inputs, yielding a total of 14 predictors for the 5-fold cross-validated multiple linear regression model. To guarantee thorough cross-validation and prevent data leakage between cross-validation folds, NMAs built in the training samples were used to extract region-specific SC-FC coupling values of the respective test samples. Prediction performance was assessed by correlating predicted and observed general intelligence scores and significance was determined with non-parametric permutation tests. NMA = Node-measure assignment. This Figure was adapted from Popp et al. (2025).

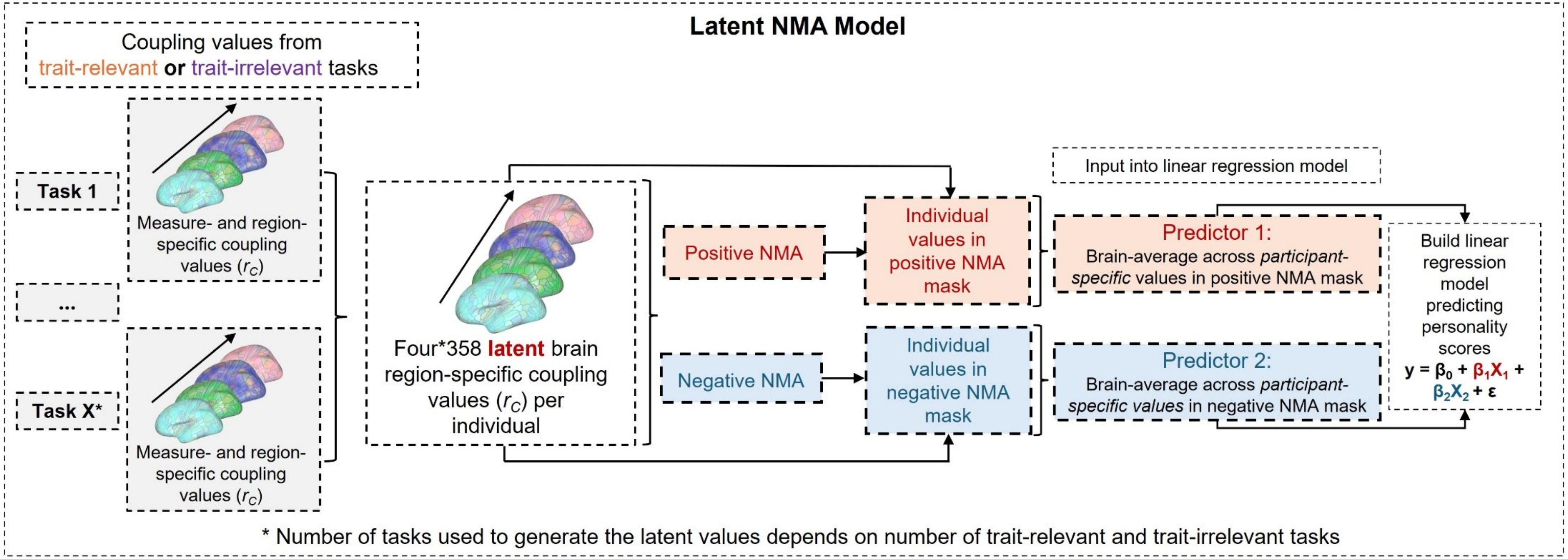


**Figure S3.** Visualization of the Latent NMA modeling approach used to predict individual personality scores from region-specific SC-FC coupling. For each personality trait, two separate prediction models were created by combining brain region-specific SC-FC coupling values from a) all trait-relevant tasks and b) all trait-irrelevant tasks into latent SC-FC coupling values before creating group-general NMA masks based on the largest positive magnitude association (positive NMA) and the largest negative magnitude association (negative NMA) between laten region-specific SC-FC coupling values and the respective personality trait. Note that latent values were either a latent variable (factor 1; if more than two tasks were determined trait-relevant) or the first component of PCA (if only two tasks were determined trait-relevant) from region-specific SC-FC coupling values of the same coupling measure. Again, brain-averages across the participant-specific values in the positive NMA mask (predictor 1) and the negative NMA mask (predictor 2) were used as model inputs for the cross-validated multiple linear regression model. To guarantee thorough cross-validation and prevent data-leakage between cross-validation folds, NMAs built in the training samples were used to extract latent region-specific SC-FC coupling values in the respective test samples. Prediction performance was assessed by correlating predicted and observed general intelligence scores and significance was determined with non-parametric permutation tests. NMA = Node-measure assignment.

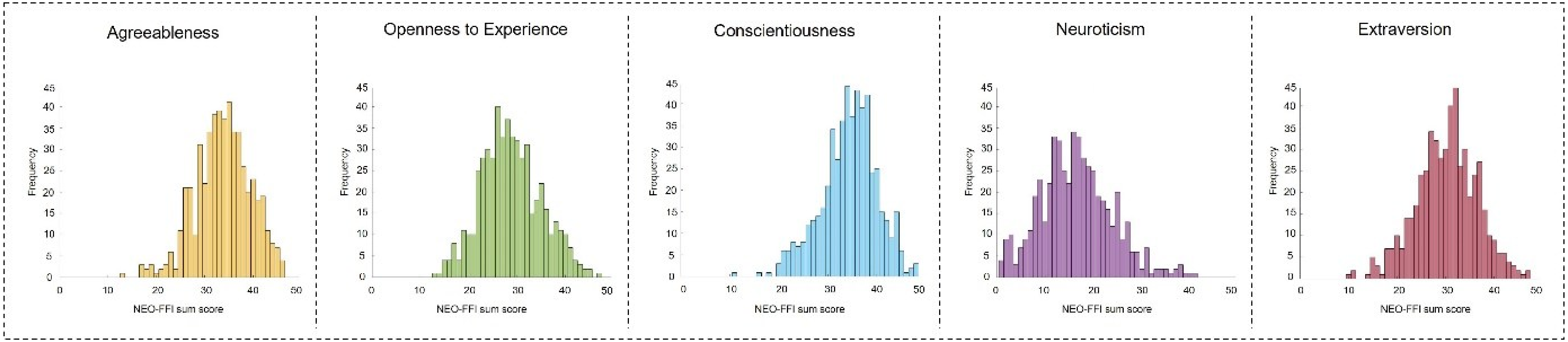


**Figure S4.** Distribution of personality scores. In the main sample (HCP; *N* = 532), agreeableness scores ranged from 13 – 46 (*M* = 33.8; *SD* = 5.7), openness to experience scores from 13 – 47 (*M* = 28.7; *SD* = 6.2), conscientiousness scores from 11 – 48 (*M* = 34.3; *SD* = 5.7), neuroticism scores from 1 – 41 (*M* = 16.5; *SD* = 7.47) and extraversion scores from 10 – 47 (*M* = 30.2; *SD* = 6.3).

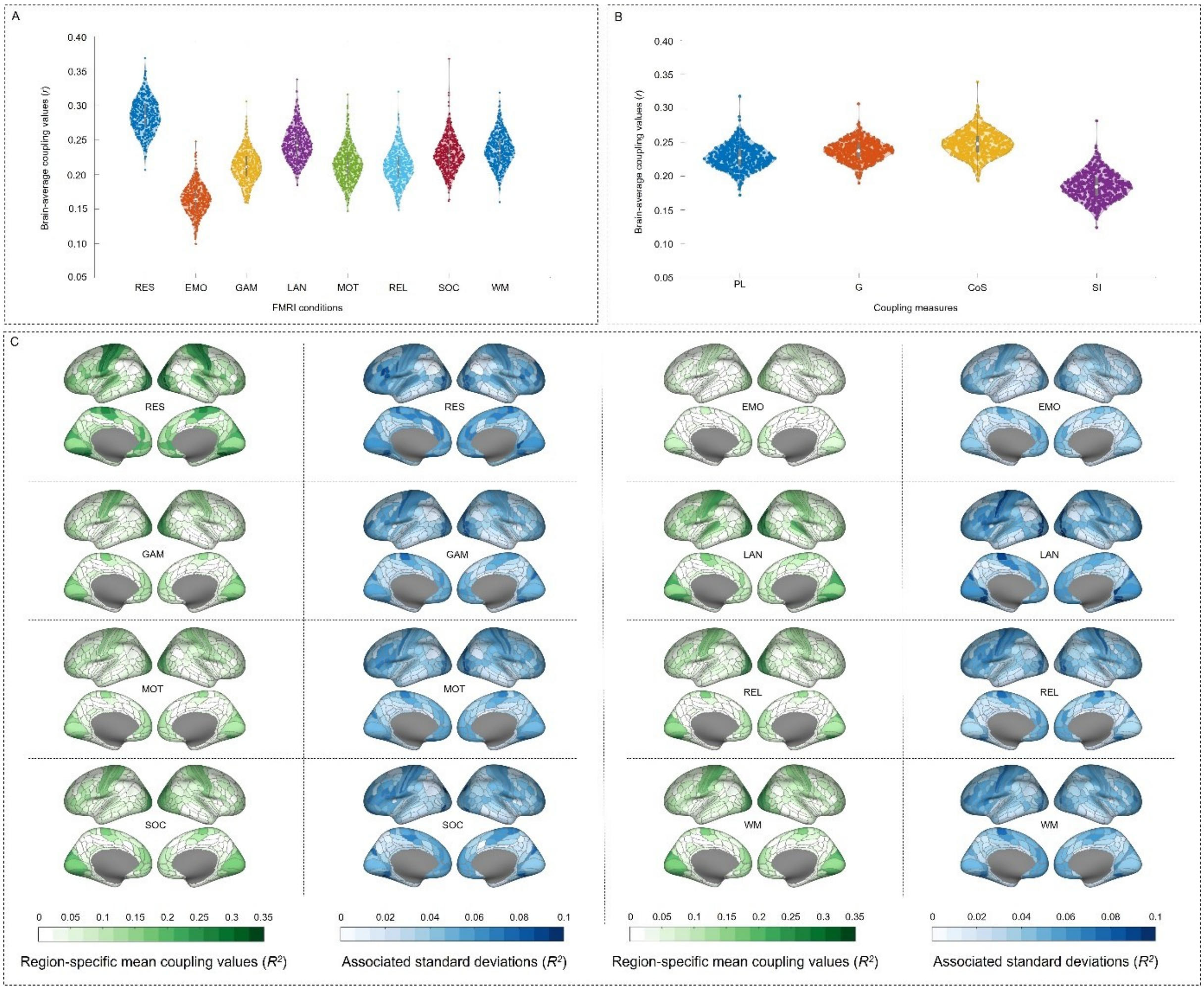


**Figure S5.** Structural-functional brain network coupling varies between conditions, coupling measures and participants. (A) Distribution of individual condition-specific brain-average SC-FC coupling values. (B) Distribution of individual measure-specific brain-average SC-FC coupling values. (C) Condition-specific group-average pattern of SC-FC coupling strength (in green). Blue maps indicate the variance in SC-FC coupling strength (as standard deviation) across participants. RES = Resting state; EMO = Emotion processing task; GAM = Gambling task; LAN = Language task; MOT = Motor task; REL = Relational processing task; SOC = Social cognition task; WM = Working memory task; PL = Path length; G = Communicability; CoS = Cosine similarity; SI = Search information. This Figure was adapted from Popp et al. (2025).

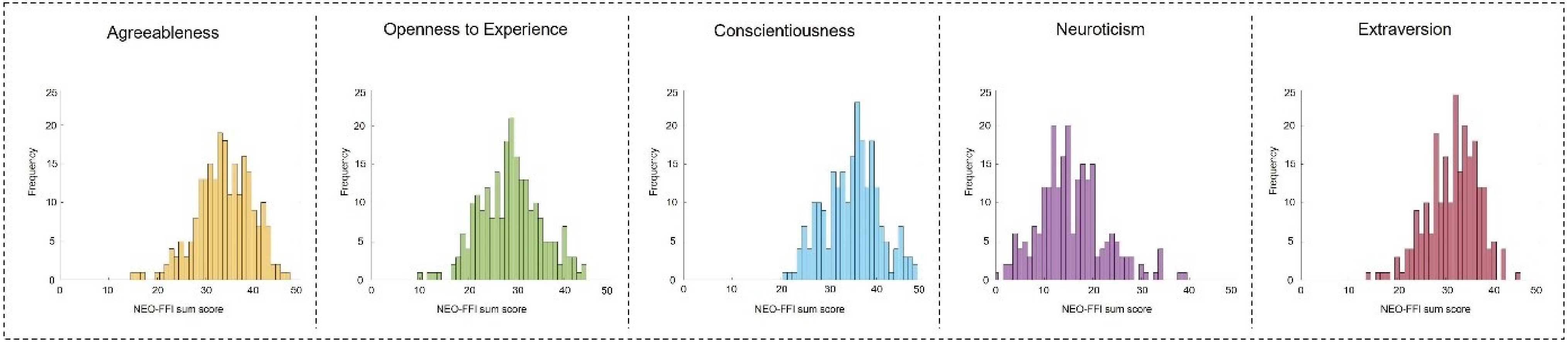


**Figure S6.** Distribution of personality scores in lockbox sample. In the lockbox sample (HCP; *N* = 232), agreeableness scores ranged from 15 – 47 (*M* = 33.9; *SD* = 5,8), openness to experience scores from 10 – 44 (*M* = 28.8; *SD* = 6.3), conscientiousness scores from 21 – 48 (*M* = 34.9; *SD* = 5.7), neuroticism scores from 0 – 39 (*M* = 15.7; *SD* = 6.9) and extraversion scores from 14 – 45 (*M* = 31.5; *SD* = 5.3).

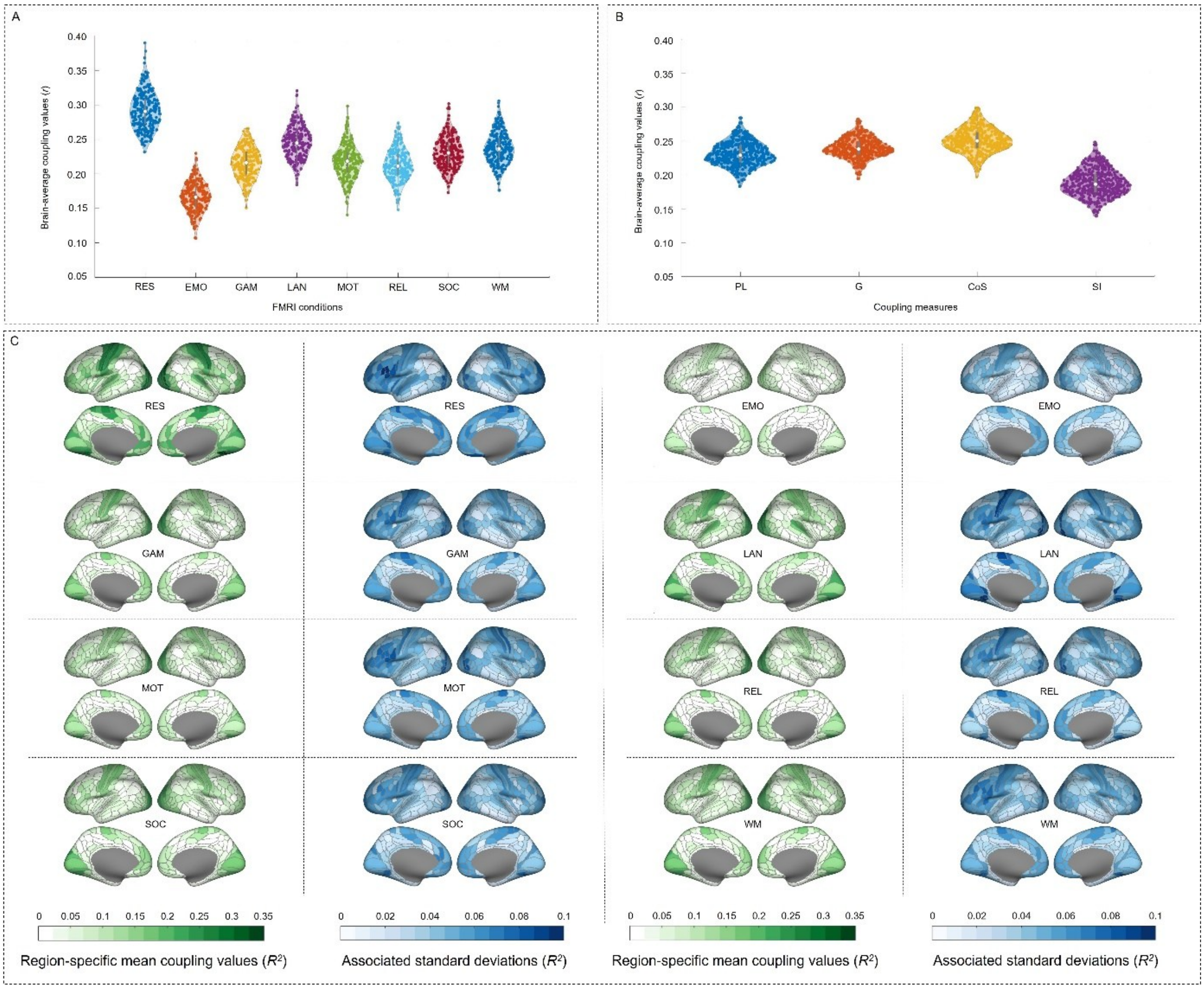


**Figure S7.** Structural-functional brain network coupling varies between conditions, coupling measures, and participants in the lockbox sample. (A) Distribution of individual condition-specific brain-average SC-FC coupling values. (B) Distribution of individual measure-specific brain-average SC-FC coupling values. (C) Condition-specific group-average pattern of SC-FC coupling strength (in green). Blue maps indicate the variance in SC-FC coupling strength (as standard deviation) across participants. RES = Resting state; EMO = Emotion processing task; GAM = Gambling task; LAN = Language task; MOT = Motor task; REL = Relational processing task; SOC = Social cognition task; WM = Working memory task; PL = Path length; G = Communicability; CoS = Cosine similarity; SI = Search information. This Figure was adapted from Popp et al. (2025).

**Supplementary Methods**

**Determination of assumed trait-relevance of tasks**

Agreeableness reflects the tendency to maintain social stability, express empathy and consider others' needs and emotions (DeYoung & Gray, 2009). Accordingly, the emotion processing task and the social cognition task were deemed trait-relevant. Openness to experience represents an individual's desire for novelty and engagement with new information. It has been linked to cognitive flexibility, attention and working memory and is positively related to intelligence (Allen & DeYoung, 2017; DeYoung, 2010). Consequently, cognitively demanding tasks - namely the language processing task, the relational processing task, and the working memory task - were considered trait-relevant. As conscientiousness encompasses top-down control of behavior and aim for achievement (DeYoung & Gray, 2009; Nostro et al., 2018), the gambling task was identified as trait-relevant. Given that neuroticism is known to be associated with an individual's sensitivity to punishment and negative affect (e.g., Allen & DeYoung, 2017; DeYoung & Gray, 2009; Nostro et al., 2018), the gambling task was considered trait-relevant. Lastly, extraversion has been linked to reward sensitivity and positive emotions (Allen & DeYoung, 2017; DeYoung, 2010; DeYoung & Gray, 2009), making the gambling task trait-relevant. Additionally, since extraversion is associated with preference for companionship and social stimulation (Costa & McCrae, 1980; DeYoung, 2015), the social cognition task was also identified as trait-relevant.

**Detailed description of similarity measures and communication measures**

Please note that this section was adapted from Popp et al. (2025).

*Similarity measures*

Similarity measures describe the conformity of regional structural connectivity profiles and are computed based on the SC matrix. They are ultimately represented in similarity matrices where each entry illustrates how the structural connections of brain region $i$ resemble the structural connections of brain region $j$. Note that the respective regional structural connectivity profiles are defined by matrix columns. For the computation of the similarity measure (cosine similarity; CoS) from each individual SC matrix, no further information about plausible communication strategies is included. Therefore, this operationalization serves as a 'baseline' measure of SC-FC coupling. Measure descriptions are adapted from Popp et al. (2024) and Zamani Esfahlani et al. (2022).

*Cosine similarity (CoS)*

Cosine similarity measures the resemblance between two brain regions' connectivity profiles (matrix columns) based on their orientation in an $N-1$ dimensional connectivity space, where $N$ is the number of brain regions, i.e., 358. We calculated the cosine similarity of the angle between two vectors $x = [x_1, \ldots, x_N]$ and $y = [y_1, \ldots, y_N]$ as $CoS_{xy} = \frac{x \cdot y}{\|x\| \cdot \|y\|}$, where vectors are region-specific connectivity profiles for every possible pair of brain regions (Han et al., 2012).

*Communication measures*

Communication measures reflect the how easy it is for two brain regions to communicate based on the underlying structural connections and a proposed signaling conceptualization (i.e., communication model like shortest path routing, navigation, random walks; see Avena-Koenigsberger et al., 2018; Seguin et al., 2023). This 'ease of communication' is subsequently represented in communication matrices. Particularly, each individual weighted SC matrix was transformed into three communication matrices representing three distinct communication measures.

*Path length (PL)*

Path length is a metric of how easily signals can be transmitted between two regions via their shortest path, as longer paths are more susceptible to noise, have longer delays in transmission and are energetically more costly (Avena-Koenigsberger et al., 2018; Rubinov & Sporns, 2010). In a network, each edge is associated with a cost $C$ (difficulty of traversing) and for weighted networks, this cost can be assessed by transforming the weight $\omega$ of each edge into a measure of length through $C = \omega^{-1}$. The shortest path between a pair of nodes (source node $s$ and target node $t$) is the sequence of edges $\pi_{s \to t} = \{A_{si}, A_{ij}, \dots, A_{mt}\}$ minimizing the sum $C_{si} + C_{ij} + \cdots + C_{mt}$ (where $C_{si}$ is the cost of traversing the edge between region $s$ and $i$) and $i, j$ and $m$ are nodes along the shortest path.

*Communicability (G)*

Communicability assumes that neural signaling unfolds as a diffusive broadcasting process, supposing that information can flow along all possible walks between two brain regions (Andreotti et al., 2014; Seguin et al., 2020). It can be characterized as the weighted sum of all walks of all lengths between two respective regions (Estrada & Hatano, 2008), where an edge is the connection between two brain regions and a walk is a sequence of traversed edges. This measure accounts for all possible connections between regions but includes walk lengths $(l_w)$ and penalizes the contribution of walks with increased lengths. For weighted networks, SC matrices $(A)$ are first normalized as $A' = D^{-1/2}AD^{-1/2}$, where $D$ is the degree diagonal matrix (Crofts and Higham, 2009). The normalized matrix is then exponentiated to calculate the communicability as $G = e^{A'}$ or $G = \sum_{wl=0}^{\infty} \frac{A'^{l_w}}{l_w!}$, where each walk is inversely proportional to its length thus 1-step walks contribute $\frac{A'^1}{1!}$, 2-step walks $\frac{A'^2}{2!}$ and so on.

*Search Information (SI)*

Search information is a measure of network navigability without global knowledge (Goñi et al., 2014; Rosvall et al., 2005). It is related to the probability that a random walker will travel between two nodes via their shortest path (i.e., the path connecting two nodes via fewest intermediate stations/nodes). This probability increases with an expanding number of paths that are available for a certain communication process to take place (Avena-Koenigsberger et al., 2018; Goñi et al., 2014). Given the shortest path between brain regions $s$ (source node) and $t$ (target node): $\pi_{s\to t} = \{s, i, j, \ldots, l, m, t\}$, the probability of accessing this shortest path is expressed as $F(\pi_{s\to t}) = f_{si} \times f_{ij} \times \ldots \times f_{lm} \times f_{mt}$, where $f_{ij} = \frac{A_{ij}}{\sum_j A_{ij}}$ and $i, j, l$ and $m$ are nodes along the shortest path. The information that is then required to access the shortest path from $s$ to $t$ is $SI(\pi_{s\to t}) = \log_2 [F(\pi_{s\to t})]$ (Goñi et al., 2014).

As search information and path length depict difficulty of communication as opposed to ease of communication, respective communication matrices were transformed to also reflect ease of communication and therefore allow easier interpretation of analysis results (for details refer to Popp et al., 2024). Further, communication matrices for search information are asymmetric, suggesting that the ease of communication between region $i$ to $j$ doesn't necessarily equal the ease of communication between region $j$ and $i$ (Seguin et al., 2019). To ensure comparability between similarity and communication measures, respective individual communication matrices were symmetrized.
